# Bacterial NLR-related proteins protect against phage

**DOI:** 10.1101/2022.07.19.500537

**Authors:** Emily M. Kibby, Amy N. Conte, A. Maxwell Burroughs, Toni A. Nagy, Jose A. Vargas, L. Aravind, Aaron T. Whiteley

## Abstract

Bacteria use a wide range of immune systems to counter phage infection. A subset of these genes share homology with components of eukaryotic immune systems, suggesting that eukaryotes horizontally acquired certain innate immune genes from bacteria. Here we show that proteins containing a NACHT module, the central feature of the animal nucleotide-binding domain and leucine-rich repeat containing gene family (NLRs), are found in bacteria and defend against phages. NACHT proteins are widespread in bacteria, provide immunity against both DNA and RNA phages, and display the characteristic C-terminal sensor, central NACHT, and N-terminal effector modules. Some bacterial NACHT proteins have domain architectures similar to human NLRs that are critical components of inflammasomes. Human disease-associated NLR mutations that cause stimulus-independent activation of the inflammasome also activate bacterial NACHT proteins, supporting a shared signaling mechanism. This work establishes that NACHT module-containing proteins are ancient mediators of innate immunity across the tree of life.

## Introduction

Bacteria are in constant conflict with viruses called bacteriophages (phages) and have evolved elaborate antiphage signaling systems to halt infections. These phage defense systems are typically multi-gene operons encoding proteins that cooperate to sense infection and inhibit virion production through diverse mechanisms (Bernheim and Sorek, 2020). The best understood antiphage systems are restriction-modification and CRISPR/Cas; however, there are many additional antiphage genes/systems we are only beginning to understand (Doron et al., 2018; Gao et al., 2020; Burroughs and Aravind, 2020; Millman et al., 2022; Rousset et al., 2022; Vassallo et al.). Most antiphage systems are a form of innate immunity, meaning they protect against a wide variety of phage and do not require previous exposure, unlike CRISPR/Cas systems which are a form of adaptive immunity. Bacteria typically encode multiple antiphage systems, often on mobile genetic elements, which are shared across the pangenome. This arsenal of antiphage systems creates a ‘pan-immune system’ (Bernheim and Sorek, 2020), which depends on the ability of antiphage genes to function well in diverse host cells, protect against disparate phages, possess potential addiction modules, and encode most of their essential components in one gene/operon.

The endeavor to catalogue antiphage signaling systems from the bacterial pan-immune system has led to an unexpected finding: some bacterial antiphage proteins are homologous to core components of the human immune system. One clear example is bacterial cyclic-oligonucleotide-based antiphage signaling systems [CBASS (Burroughs and Aravind, 2020; Cohen et al., 2019; Morehouse et al., 2020; Whiteley et al., 2019)], which are homologous to the human cGAS and STING proteins. Other examples are bacterial Viperins (Bernheim et al., 2021) and bacterial Gasdermins (Johnson et al., 2022), which are homologous to human Viperin and Gasdermin, respectively. These genes are antiviral in both humans and bacteria, and bioinformatic evidence supports that these genes share a common ancestor (Bernheim et al., 2021; Burroughs and Aravind, 2020; Burroughs et al., 2015; Johnson et al., 2022). It appears that the pervasive horizontal gene transfer of antiphage systems between bacteria may have also resulted in metazoans horizontally acquiring antiphage genes from bacteria, which were then adapted to fight viruses in eukaryotic cells. We therefore hypothesized that additional components of the metazoan innate immune system originated from antiphage signaling systems and searched for those genes in bacteria.

## Results

### A bacterial NACHT protein is antiphage

Antiphage systems frequently cluster together into “defense islands” throughout bacterial genomes (Makarova et al., 2011). This phenomenon has been used to identify novel antiphage systems by interrogating genes of unknown function that are colocated with known antiphage systems (Doron et al., 2018; Gao et al., 2020; Millman et al., 2022). We investigated genes of unknown function located near CBASS systems and identified a gene encoding a NACHT protein module in *Klebsiella pneumoniae* MGH 35 (**Figure 1A and B**, WP_015632533.1). We named this gene bacterial NACHT module-containing protein 1 (bNACHT01), and it did not appear to be in an operon with any other genes. We hypothesized bNACHT01 was antiphage and tested that hypothesis by expressing bNACHT01 from its endogenous promoter in *E. coli*, then challenging those bacteria with diverse phages. bNACHT01 conferred over a 100-fold increase in protection against phage T4 and over a 1000-fold increase in protection against phages T5 and T6 (**Figure 1C and D**). Within an hour post-infection, the phage lysed cultures of bacteria expressing an empty vector while bacteria expressing bNACHT01 were protected (**Figure 1E**).

**Figure 1.**
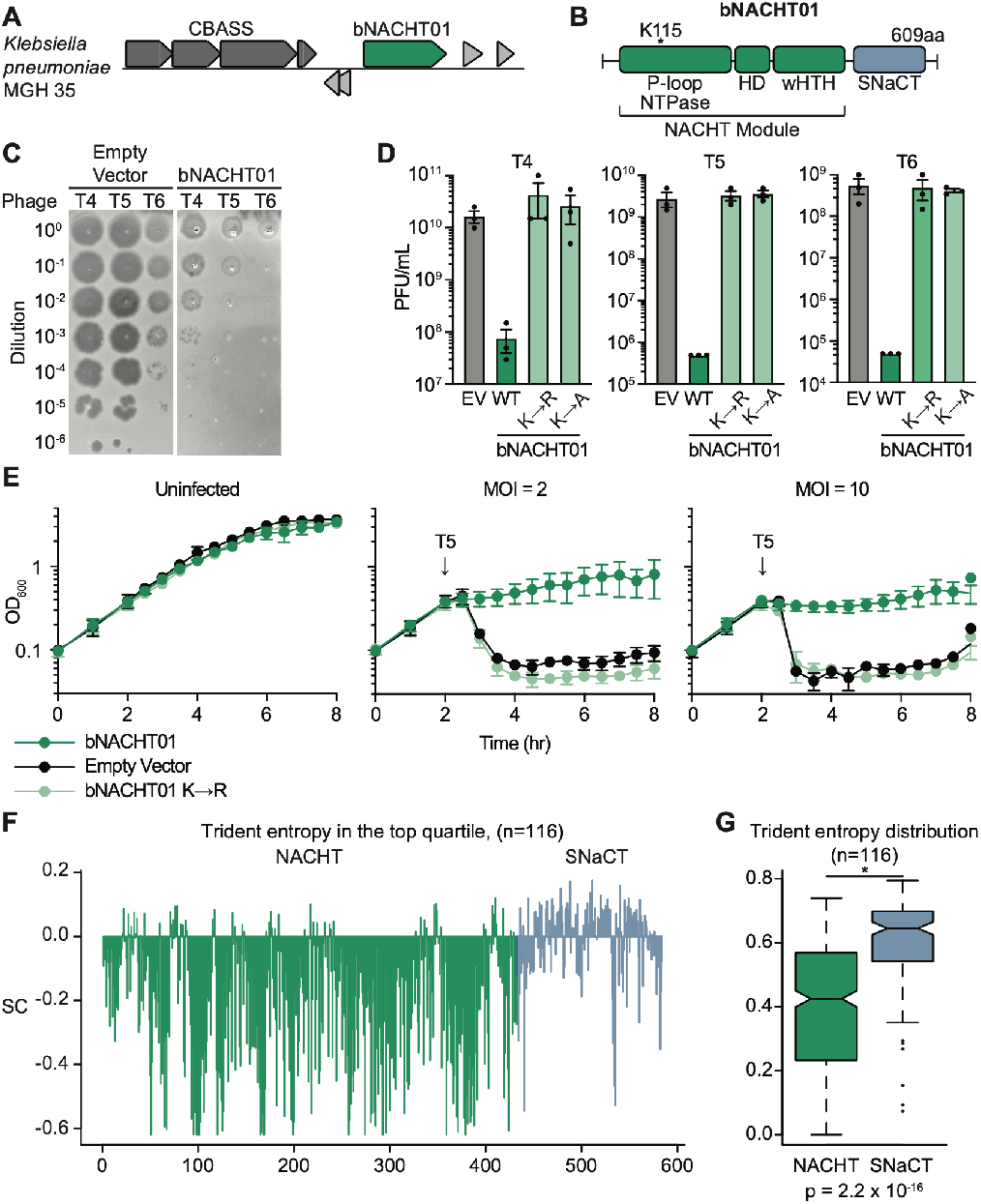
A bacterial NACHT module-containing protein is antiphage. **(A)** Genome context of *bNACHT01*, which is located near a CBASS system in *Klebsiella pneumoniae* MGH 35. **(B)** Schematic of bNACHT01 (WP_015632533.1) protein domains, annotated by alignment to the NACHT module of NLRC4. The P-loop NTPase domain is also known as a nucleotide-binding domain (NBD), the helical domain (HD), and the winged helix-turn-helix (wHTH, also called WHD for winged helical domain) are indicated. See **Figure S1** for a protein alignment of bNACHT01 with eukaryotic NACHT modules. **(C)** Efficiency of plating of indicated phages infecting *E. coli* expressing bNACHT01 or an empty vector (EV). Data are representative images of *n = 3* biological replicates. **(D)** Efficiency of plating of phages infecting *E. coli* expressing the indicated genotype. Data represent the mean ± standard error of the mean (s.e.m.) of *n = 3* biological replicates, shown as individual points. **(E)** Growth curve of *E. coli* expressing the indicated plasmid. Arrows indicate the time each culture was infected with phage T5 at the indicated multiplicity of infection (MOI). **(F)** Scaled Trident entropy (SC) values for individual residues of bNACHT01-like proteins. The trident entropy for each column of the alignment, including both the NACHT module and SNaCT domain, is scaled with respect to the top quartile. Positions with values greater than 0 are those with diversity in the top quartile. **(G)** Distribution of trident entropy (S) values across NACHT and SNaCT modules in bNACHT01-like proteins showing significantly different mean Trident entropy. Values were compared using a two-sample t-test.

The NACHT module of bNACHT01 shares core features with NACHT modules in eukaryotes, including a Walker A motif that binds NTPs in those proteins (Hu et al., 2013; Kim et al., 2016; Sandall et al., 2020) (**Figure 1B** and **Figure S1**). Mutation of the conserved lysine residue within the bNACHT01 Walker A motif (K115) to arginine or alanine abrogated phage defense (**Figure 1D and E** and **Figure S1**) indicating that NTP binding is required for antiviral function.

We next interrogated the multiple sequence alignment of the clade of bacterial NACHTs defined by bNACHT01 to better understand the mechanism of phage defense. This analysis revealed that the NACHT module is relatively stable in its sequence conservation; however, the region to the C-terminus of the NACHT module appears to be rapidly diversifying (**Figure 1F and G).** We named this region the **<u>S</u>**hort **<u>N</u>**ACHT-**<u>a</u>**ssociated **<u>C</u>**-**<u>T</u>**erminal domain (SNaCT). We used trident entropy scores to compare the degree of amino acid conservation for the NACHT and SNaCT domains and confirmed that the mean entropy score is higher for SNaCT domain (**Figure 1F and G**) (Valdar, 2002). The rapid diversification of the SNaCT domain is a hallmark of a host-pathogen “arms race”, where evolutionary pressure from interactions between an immune sensor and pathogen selects for amino acid substitutions that change protein-protein interactions. In eukaryotic NACHT proteins, the C-terminus is often the “sensor” or “receptor” region of the protein that responds to infection stimuli (Tenthorey et al., 2017). It is therefore paradoxical that bNACHT01 is both capable of detecting a variety of unrelated phages (T4, T6, and T5) and rapidly diversifying in the sensor region. Instead, these data might suggest that bNACHT01 SNaCT domain is evolutionarily diverging under pressure from constant antagonism by phage-encoded proteins that enable immune evasion (e.g., phage encoded bNACHT inhibitors).

### Diversity and ubiquity of NACHT module-containing proteins in bacteria

The NACHT protein module was first discovered in eukaryotes where it is often found in proteins that mediate immunity and inflammation. The best understood metazoan NACHT proteins belong to the nucleotide-binding domain and leucine-rich repeat-containing gene family (NLRs) (Ting et al., 2008), which have a core nucleotide-binding and oligomerization domain (NOD) whose function is fulfilled by a NACHT module. Mammalian NLRs are immune components that play an important role in the formation of inflammasomes (such as NAIP/NLRC4, NLRP1, and NLRP3), transcriptional regulation (CIITA), and other inflammatory responses (Sandall et al., 2020). NACHT modules are also found in Fungi where HetE/D proteins can mediate self-nonself discrimination after two hyphal cells have fused their cytosols in a process called heterokaryon incompatibility (Dyrka et al., 2014; Saur et al., 2021; Uehling et al., 2017).

NACHT proteins have been rigorously investigated in eukaryotes, but little is known about their roles in bacteria. The NACHT protein module belongs to the large family of STAND NTPases, which describes many divergent proteins. Both active and inactive (with disrupted Walker A/B motifs) STAND NTPase domains were previously identified computationally in predicted antiviral conflict systems that are enriched in multicellular bacteria (Kaur et al., 2020, 2021). STAND NTPases were also observed in the Antiviral ATPase/NTPase of the STAND superfamily (AVAST) antiphage signaling system (Gao et al., 2020), as well as in prophage-encoded antiphage systems (Rousset et al., 2022), but NACHT domains were not specifically recognized in either study.

We undertook an exhaustive bioinformatic analysis of NACHT module-containing proteins across publicly available genomes of both eukaryotes and prokaryotes. We first started with identifying NACHT modules based on amino acid sequence and predicted structural features. The NACHT protein module belongs to the STAND-Cdc6-Orc family of AAA+ NTPases, which in turn belong to the ASCE division of P-loop NTPases. STAND NTPase modules are unified structurally by a characteristic C-terminal extension to the core AAA+ domain in the form of winged helix-turn-helix (wHTH) domain, also referred to as a Winged Helix Domain (WHD), which they share with Cdc6-Orc AAA+ and transposase ATPases (Koonin and Aravind, 2000; Leipe et al., 2004; Zhang et al., 2016). Within the STAND clade, the NACHT subclade is unambiguously separated from other STANDs based on characteristic motifs, including the D[GAS]hDE signature (a small amino acid directly C-terminal to the Mg^2+^-coordinating aspartate within the Walker B motif, followed by two acidic residues one position away), as well as signatures in the region N-terminal to the NTPase domain and the above-mentioned C-terminal wHTH extension (**Figure S1**) (Koonin and Aravind, 2000; Leipe et al., 2004).

Our analysis identified approximately 20,000 unique bacterial NACHT proteins. They are encoded by about 9-10% of complete, published bacterial genomes (**Table S1**). NACHT proteins are found throughout the bacterial superkingdom, including in the genomes of pathogenic bacteria, members of the human gut microbiome, and other important bacteria from environmental niches. Some bacterial phyla show a much higher tendency than average to encode NACHT proteins (**Figure S2**): we found that 58% of the cyanobacteria, 25% of the actinobacteria and 24% of the deltaproteobacteria encode NACHT proteins (**Figure S2D**). Cyanobacteria also tend to display a large number of paralogous versions per genome – for instance, a record number of 23 paralogous NACHT proteins are seen in Rivularia sp. PCC 7116 (**Figure S2A**). Moreover, organisms with 3 or more NACHT proteins have a significant tendency to be multicellular bacteria (**Figure S2B**). Notably, the multiple copies of the NACHT proteins in these organisms tend to possess distinct effector and sensor domains, suggesting that they are not merely duplications representing iterations of the same theme, but a diversified biochemical repertoire potentially optimized to deal with the unique immune challenges related to multicellularity. This situation mirrors the previously described class of immunity and apoptosis mechanisms shared by a range of multicellular bacteria (Kaur et al., 2020, 2021).

The NACHT modules from each of these proteins were aligned and related proteins were grouped into clades (**Figure 2** and **Table S1**). By aligning proteins based on the NACHT module, this analysis was independent of fused protein domains on each polypeptide (**Figure 2**). Our analysis also included NACHT module-containing proteins from Archaea and Eukaryotes, which allowed us to group related proteins from different domains of life into a total of 25 different clades and establish evolutionary relationships (**Figure 2**).

**Figure 2.**
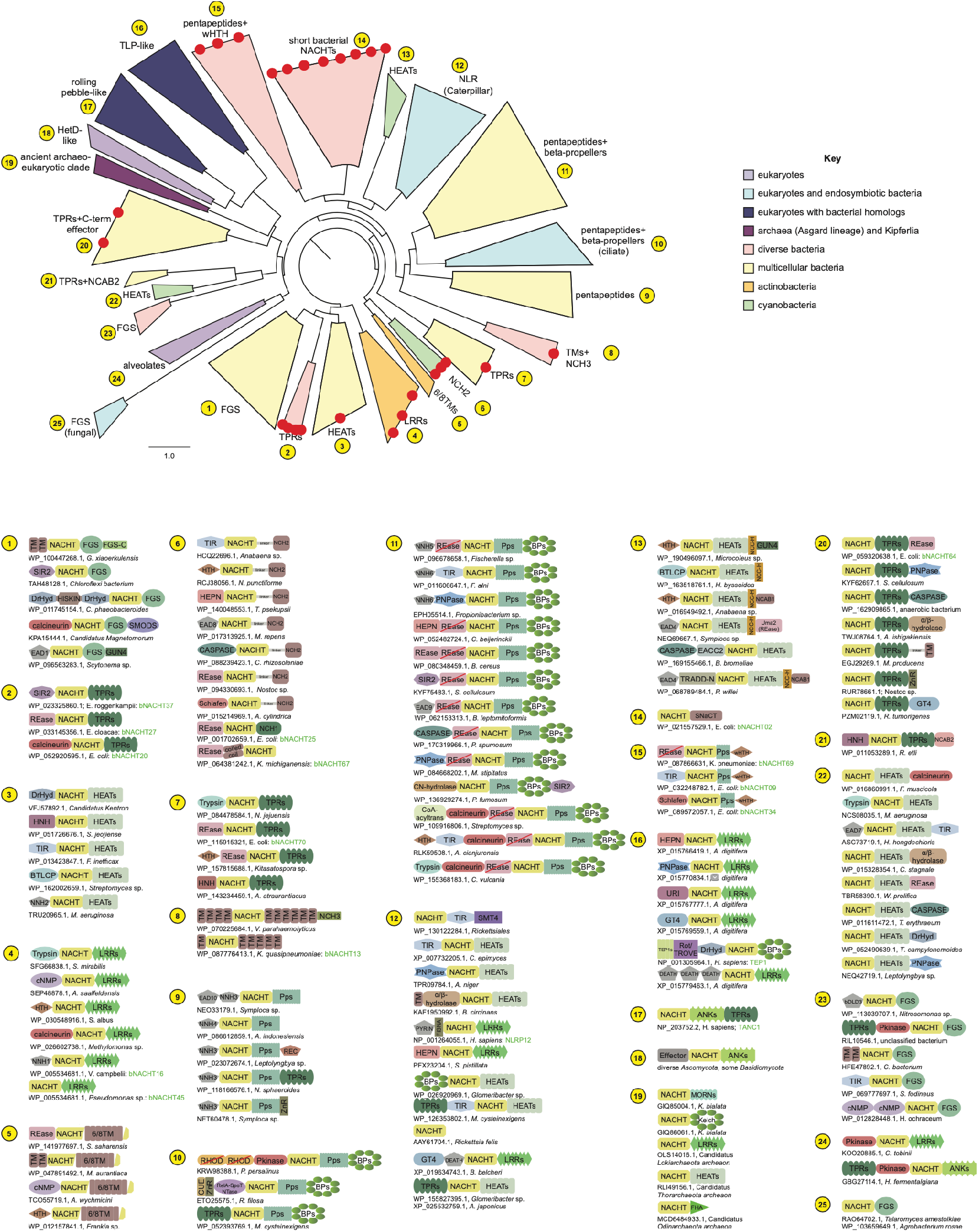
NACHT module containing proteins in bacteria are widespread and diverse. A sequence-based phylogenetic tree of NACHT modules was generated using NACHT module containing proteins from eukaryotes and prokaryotes. The NACHT module, not accessory domains, were used for tree building. Clades are color-coded based on the indicated key and numbered arbitrarily in yellow dots. Representative domain architectures for each clade are provided, including NCBI Protein accession number, species of origin, and gene name in green where appropriate. Red dots indicate the bacterial NACHT proteins from each clade that were selected for analysis in this study. 3’-5’ cyclic nucleotide-generating cyclase (cNMP), Ankyrin repeats (ANKs), Bacterial Death-like domain-3 (bDLD3), Beta-propeller repeats (BPs), bacterial transglutaminase-like cysteine protease (BTCLP), Carbon-nitrogen hydrolase (CN-hydrolase), CoA-dependent acyltransferase (CoA-acyltrans), deoxyribohydrolase (DrHyd), Effector associated constant component (EACC), Effector-associated domain (EAD), Formylglycine-generating enzyme sulfatase (FGS), FGS C-terminal domain (FGS-C), forkhead-associated domain (FHA), Fish-specific NACHT-associated domain (FISNA), Glycosyltransferase 4 (GT4), Huntington, Elongation Factor 3, PR65/A, TOR (HEAT), Higher eukaryotes and prokaryotes nucleotide binding domain (HEPN), homing endonuclease (HNH), Helix-turn-helix (HTH), Leucine rich repeat (LRR), Membrane occupation and recognition nexus (MORN),, NACHT N-terminal helical domain (NNH), NACHT C-terminal helical domain (NCH), NACHT C-terminal α / β domain (NCAB), NACHT C-terminal cysteine and histidine-containing domain (NCC-H), Protein kinase domain (Pkinase), Polyribonucleotide nucleotidyl transferase (PNPase), Pentapeptide repeat (Pp), Restriction endonuclease (REase), Receiver domain (Rec), Rhodanese domain (RHOD), Sirtuin (SIR2), Second messenger oligonucleotide or dinucleotide synthetase domain (SMODS), Telomerase associated protein 1 (TEP1), Toll/interleukin receptor (TIR), Transmembrane (TM) number indicates how many copies of this motif, Tetratricopeptide repeat (TPR), N-terminal domain of tumor necrosis factor receptor type 1 associated death domain protein (TRADD-N), UvRC and Intron-encoded endonuclease domain (URI), Zinc ribbon (ZnR). Red slash indicates catalytically inactive domain. See **Table S1** for additional details on genes used to construct the phylogenetic tree.

The C-terminal regions of bacterial NACHT proteins can be placed into different, broad categories: the antigen receptor-type, those that have transmembrane (TM) domains, those with short C-terminal extensions, or those with a combination of these features (**Table S1** and Supplementary Discussion). There are two types of antigen receptor-type domains: FGS (Formylglycine-generating enzyme sulfatase) domains and supersecondary structure-forming tandem repeats {e.g., LRR (Leucine Rich Repeat), TPR (Tetratricopeptide Repeat), HEAT (Huntington, Elongation Factor 3, PR65/A, TOR)}. Antigen receptor-type and TM domains are found across many different clades, however, bacterial NACHT proteins with short C-terminal extensions that lack supersecondary structure forming elements, such as bNACHT01, are predominantly found as the monophyletic clade 14 (**Figure 2**). These and other characteristics that predominate each NACHT clade are annotated on **Figure 2**.

The N-terminal regions of bacterial NACHT proteins encode many enzymatic domains that have previously been associated with biological conflict, including nucleases (RNases and DNases), peptidases, and NAD^+^-targeting enzymes (TIR and Sirtuin) (Aravind et al., 2022). Other domains also include Effector Associated Domains, Death-like domains, RNA-binding domains, transcription regulatory domains, and nucleotide signal-generating or degrading domains (**Table S1** and Supplementary Discussion).

The tripartite domain architecture we observe in bacterial NACHT proteins is consistent with the domain architectures previously observed in eukaryotic NACHT proteins (Koonin and Aravind, 2000; Leipe et al., 2004). The central NACHT in mammalian NLRs is flanked by an N-terminal “effector” domain that coordinates signaling and a C-terminal “sensor” domain that often consists of supersecondary structure-forming tandem repeats, such as LRRs. The similarity of the tripartite domain architecture in bacterial and mammalian NACHT proteins suggest that the sensor and effector domains of canonical eukaryotic NLRs represents a broader organizational strategy across the tree of life (Leipe et al., 2004). These observations further imply that the role of the NACHT module is to act as a signaling hub that transduces detection of an invader signal into diverse biochemical outputs, enabling the host to respond to a threat.

### Evolutionary history of NACHT proteins

Phylogenetic relationships between prokaryotic and eukaryotic NACHT proteins suggest these genes have horizontally transferred from prokaryotes to eukaryotes on multiple occasions. The clades in **Figure 2** are categorized by the organisms that are most represented in that clade. This includes eukaryotes, a mixture of eukaryotes and bacteria, as well as various groups of prokaryotic organisms. Eukaryotic NACHT modules are found in multiple clades (**Figure 2**), suggesting that NACHT modules were acquired on several distinct occasions and subsequently experienced lineage-specific expansions (Leipe et al., 2004). Fungi have acquired NACHT proteins from multiple horizontal gene transfer events and one of these resulted in the expansion of the heterokaryon incompatibility NACHT proteins (Clade 18, HetD/E-like). In the mammalian lineage, there are two distinct clades of NACHT modules. The first of these is TEP1 (previous named TP-1), found in clade 16, and was acquired early in eukaryotic evolution from bacteria. The second is typified by the mammalian NLR/Caterpillar NACHT proteins, found in clade 12, and represents a separate transfer from bacteria.

Despite these postulated horizontal gene transfer events being ancient, Clade 12 includes extant prokaryotic NACHT proteins encoded by *Rickettsiales*. The *Rickettsiales* are an order of obligate intracellular bacteria that have coevolved extensively with animals. Similarly, clade 18 includes homologs of heterokaryon incompatibility NACHTs from bacterial endosymbionts of fungi. Thus, our phylogenetic analysis suggests that metazoan and fungal hosts acquired their NACHT genes involved in immune mechanisms from obligate intracellular symbionts/pathogens. This likely origin of metazoan and fungal NLRs and NLR-related proteins, from intracellular bacteria, stands in contrast to the horizontal transfer of STING from an extracellular bacterial symbiont (Burroughs and Aravind, 2020; Morehouse et al., 2020).

### Multiple bacterial NACHT proteins provide a broad spectrum of antiphage immunity

The bNACHT01 protein was potently antiphage (**Figure 1**); however, our bioinformatic analysis demonstrated that there are many additional clades of unrelated NACHT proteins in bacteria (**Figure 2**). To measure the breadth of antiphage signaling of NACHT proteins, we expressed 27 representative NACHT module-encoding genes in our *E. coli*-based phage resistance assay (**Figures 3**, **S4**, and **S5**). Representatives were selected based on protein domain, similarity of domain architecture to eukaryotic NACHT proteins, and phylogenetic distance of the source genome to *E. coli* (to recapitulate native host cell conditions). Bacterial NACHT proteins were expressed from promoters in their native genomic context and were only rarely within poorly conserved operons (**Table S2**). Bacteria were challenged with a diverse panel of double stranded DNA (T2–T7, λvir), single-stranded DNA (M13), and positive-sense single stranded RNA (MS2, Qβ) phages. We also included a previously characterized CBASS system from *Vibrio cholerae* and a restriction modification system from *E. coli* as positive controls in these experiments. Diverse bacterial NACHT proteins from different clades exhibited robust antiphage signaling across a wide range of phages (**Figures 3**, **S4**, and **S5**). Intriguingly, some bacterial NACHT proteins defended against both DNA and RNA phages. These data are the first example of a bacterial innate immune system that defends against an RNA phage.

**Figure 3.**
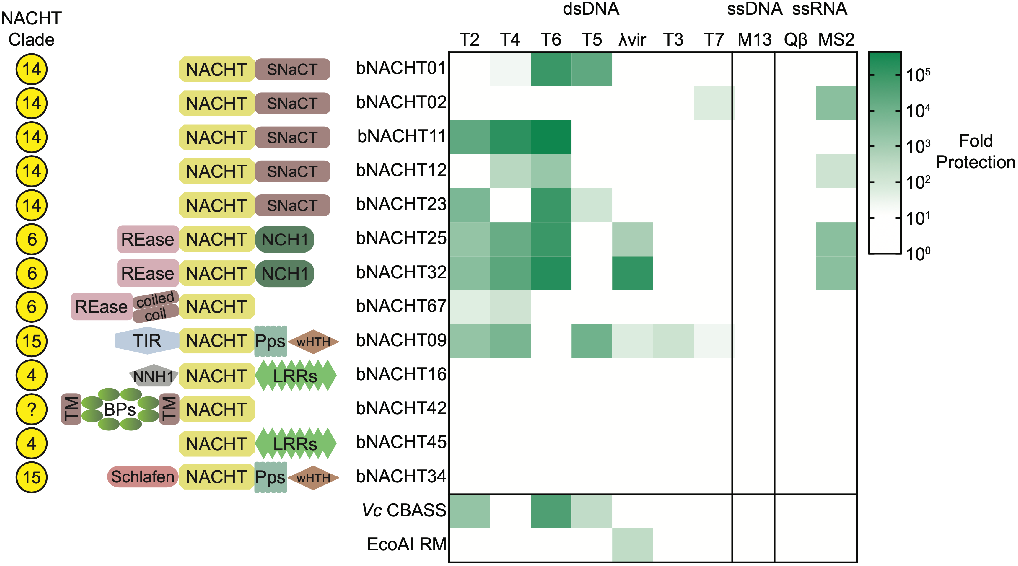
Bacterial NACHT proteins are antiphage. Heat map of fold defense provided by the indicated bNACHT gene for a panel of diverse phages. *E. coli* expressing the indicated defense system was challenged with phages and fold defense was calculated for each defense system-phage pair by dividing the efficiency of plating (in PFU/mL) on empty vector by efficiency of plating on defense system-expressing bacteria. The NACHT clade and domain architecture for each bNACHT are shown. bNACHT genes displayed in this figure are a subset of the 27 interrogated, selected based on their robust antiphage activity or the diversity of domain architectures sampled. *Vibrio cholera* CBASS (*Vc* CBASS) and *E. coli* UPEC-36 restriction modification system (EcoAI RM) were included as positive controls. Data represent the mean of *n* = *3* biological replicates. Domain abbreviations as described in **Figure 2**. See **Table S2** and **Figures S3** and **S4** for details on all 27 bNACHT genes analyzed. See **Figure S5** for raw efficiency of plating data.

Our interrogation of a wide range of bacterial NACHT proteins demonstrates that bacterial NACHT proteins are related to metazoan NLRs because they share a number of features: (1) a broad role in antipathogen immune signaling, (2) conserved sequence and structural characteristics of their NACHT modules, and (3) a characteristic tripartite protein architecture (C-terminal tandem repeat domains, central NACHT, and N-terminal effector domain). These findings show that role of STAND NTPases, including the bacterial NACHTs characterized herein, extend the scope of STAND NTPases in antiviral defense beyond the previously proposed AVAST systems (Gao et al., 2020).

### Phage proteins alter bacterial NLR signaling

We sought to understand how phages alter bacterial NACHT protein signaling by generating phage mutants that evade defense signaling (suppressor mutant phage). Phage T5 was selected for analysis. Wild-type T5 plaque formation is robustly inhibited by bNACHT01 (**Figure 1D and E**); however, when bacteria were infected with a high number of T5 PFU, suppressor mutants capable of escaping bNACHT01 and forming a plaque were isolated (**Figure 4A**). These mutants were extremely rare, appearing at a rate of one suppressor for every 5 x 10^7^ PFU of wild-type phage (**Table S3**). Accordingly, genome sequencing revealed that nearly every suppressor phage encoded at least two mutations that affected the same ORFs: one mutation that altered *orf008* (accession number AAX11945.1) and the other that altered *orf015* (accession number AAX11952.1, **Tables S4** and **S5**). The majority of the suppressor mutations identified were missense mutations, although some included frameshifts, nonsense, and promoter mutations (**Tables S4** and **S5**). Genes *orf008* and *orf015* are encoded in the “first-step transfer” region of the T5 genome, an 11 kb section that is injected first into host bacteria and coordinates injection pausing for approximately five minutes before the remainder of the genome is injected (Davison, 2019). During those first minutes of infection, other pre-early genes of the first-step transfer region remodel core processes and shut down signaling within the host cell (Davison, 2015).

**Figure 4.**
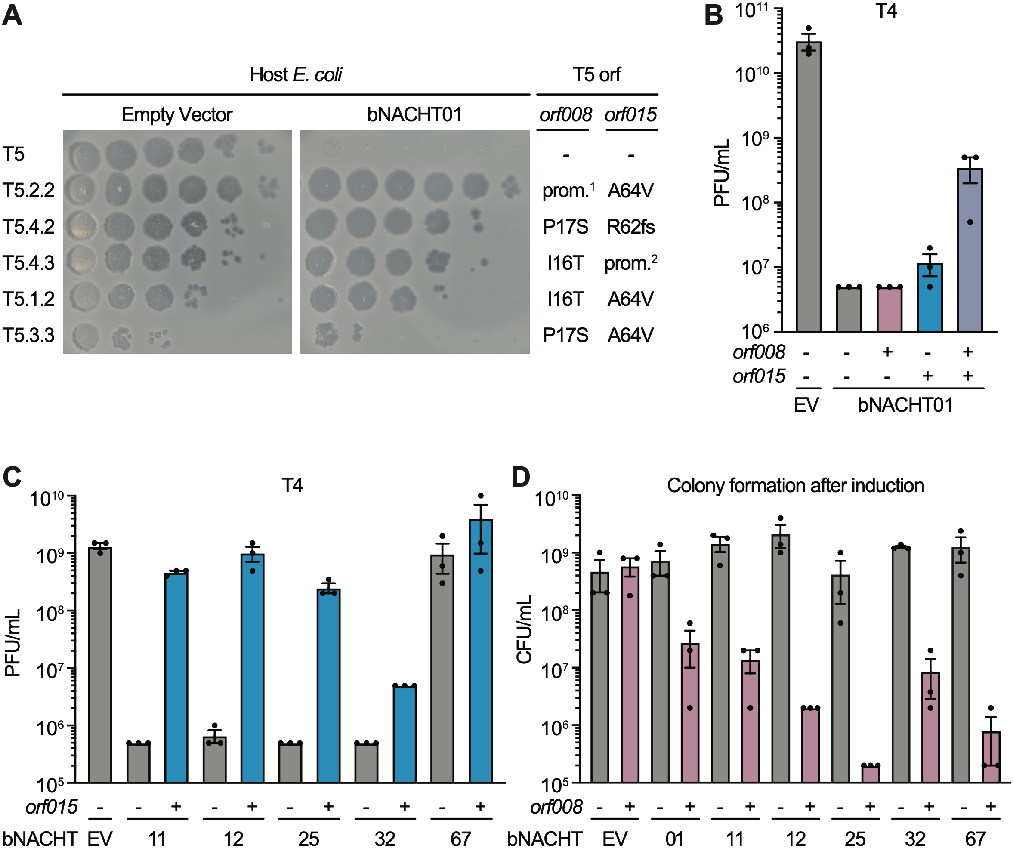
Phage proteins modulate host immune responses. **(A)** Efficiency of plating of wild-type or suppressor T5 phage when infecting *E. coli* expressing the indicated plasmid. Impact of *orf008* and *orf015* suppressor mutations are indicated. Data are representative images of *n = 3* biological replicates. Wild-type alleles (-); mutations in the promoter region of *orf008* (prom^1^); frame shift, mutations at the indicated position (fs); and mutations deleting *orf009–012* predicted to disrupt the promoter of *orf015* (prom^2^) are indicated. See **Table S3** for rates of suppressor phage isolation, **Table S4** for suppressor mutations identified in *orf008* and *orf015*, and **Table S5** for a complete list of mutations identified. **(B)** Efficiency of plating of phage T4 when infecting *E. coli* expressing bNACHT01 or an empty vector (EV) on one plasmid and the indicated phage T5 gene(s) on a second plasmid. See **Figure S5** for efficiency of plating of phage T6. **(C)** Efficiency of plating of phage T4 when infecting *E. coli* expressing the indicated bNACHT gene on one plasmid and phage T5 *orf015* on a second plasmid. See **Figure S6** for efficiency of plating of phages T2 and T6. **(D)** Quantification of colony formation of *E. coli* expressing the indicated bNACHT system on one plasmid and *orf008* on a second plasmid. See **Figure S7** for colony formation in the presence of *orf015*. For B–D, expression of *orf008, orf015*, and *mCherry* is IPTG-inducible. (-) symbols denote induction of an *mCherry* negative control. (+) symbols denote induction of the indicated phage gene. Data represent the mean ± s.e.m. of *n* = *3* biological replicates, shown as individual points.

We hypothesized that the low frequency of isolating suppressor phages reflect that T5 must encode mutations in both *orf008* and *orf015* to evade bNACHT01 signaling. To measure the impact of these genes on bNACHT01 antiphage signaling, we constructed an assay where bNACHT01 was co-expressed with either *orf008*, *orf015*, or both phage genes, then challenged with phages T4 and T6 (**Figure 4B** and **Figure S6**). bNACHT01 provided 1000-fold protection against phage T4 in this assay, and expression of either *orf008* or *orf015* individually had a modest impact on the efficiency of plating. However, expression of both genes together resulted in a 100-fold recovery of phage T4 virulence (**Figure 4B**). These data suggest that *orf008* and *orf015* act together to allow phage to evade bNACHT01 signaling.

Relatively few bacterial NACHT proteins protected *E. coli* against phage T5 (**Figure 3** and **Figure S4**), and we hypothesized that *orf008* and *orf015* might be broadly responsible for T5 immune evasion. To test this, we selected bNACHT alleles that defended against phage T4, but not phage T5, then repeated our assay for measuring the impact of these phage genes. Expression of *orf015* significantly decreased the protection by bNACHT11, 12, 25, and 32 against phage T4 (**Figure 4C**). Similar results could be obtained for a subset of these genes when phage T2 and T6 were used (**Figure S7**). We next analyzed the effect of *orf008* on bNACHT signaling. Interestingly, we found that bacterial growth was inhibited when *orf008* was co-expressed with bNACHT alleles but not when co-expressed with empty vector (**Figure 4D**). The growth inhibition was specific to *orf008*; *orf015* did not alter bacterial growth when expressed with bNACHT alleles (**Figure S8**). Because *orf008* inhibited growth, we did not measure its impact on phage defense beyond bNACHT01 (**Figure 4B**). Taken together, these data demonstrate that two phage genes alter signaling of a wide variety of bNACHT alleles and provide evidence for a complicated relationship between phage genes and bNACHT host defense systems.

### Human disease mutations activate bacterial NLRs

Human NLR protein activation has potent signaling consequences and rare, monoallelic mutations in patients cause serious diseases that include bare lymphocyte syndrome, Crohn’s/inflammatory bowel disease, and autoimmune conditions (Kim et al., 2016; Zhong et al., 2013). A subset of these diseases are inflammasomopathies, which are point mutations in NLRs that result in stimulus independent hyperactivation of inflammasome signaling. Patients encoding H443P mutations in NLRC4 display familial cold autoinflammatory syndrome (FCAS) (Kitamura et al., 2014) and H443L mutations also result in NLRC4 activation in cells (Hu et al., 2013). Histidine 443 is a highly conserved and defining residue located within the wHTH (WHD) domain of the NACHT module (**Figure S9**). In NLRC4 H443 is thought to interact with ADP to stabilize an inactive conformation (Hu et al., 2013).

Given the high degree of conservation between human and bacterial NACHT modules, we hypothesized that mutations which hyperactivate human NLRs might also hyperactivate bacterial NLR-related proteins. Structure-guided alignments between NLRC4 and bNACHT25 identified the analogous residue to H443 and mutations at this location were constructed in bNACHT25 (**Figure S9**). The effector domain of bNACHT25 is a predicted Mrr-like restriction endonuclease (REase). When activated, bNACHT25 is predicted to cleave DNA, resulting in toxicity to the host cell and/or destruction of the DNA phage chromosome (**Figure 3**, **S3**, and **Table S2**). For this reason, we expressed wild-type bNACHT25 and mutant alleles using an inducible system. Expression of the histidine mutant, but not wild-type bNACHT25, resulted in potent bacterial growth inhibition (**Figure 5A**). An additional mutation predicted to disrupt the nuclease activity of the effector domain (D48A) rescued the growth inhibition of hyperactive bNACHT25 (**Figure 5A**). We next interrogated bNACHT16 because this gene is highly similar to human NLR proteins, encoding leucine-rich repeats (LRRs) at the C-terminus (**Figure 3**, **Figure S3**, **Table S2**). Introduction of mutations at the H443 equivalent residue of bNACHT16 also resulted in bacterial growth inhibition, consistent with NLR hyperactivation (**Figure 5B**). Histidine to leucine mutations, shown to synthetically activate NLRC4 (Hu et al., 2013), or histidine to proline mutations, found in patients with inflammasomeopathies (Kitamura et al., 2014), both inhibited growth equivalently (**Figure 5**).

**Figure 5.**
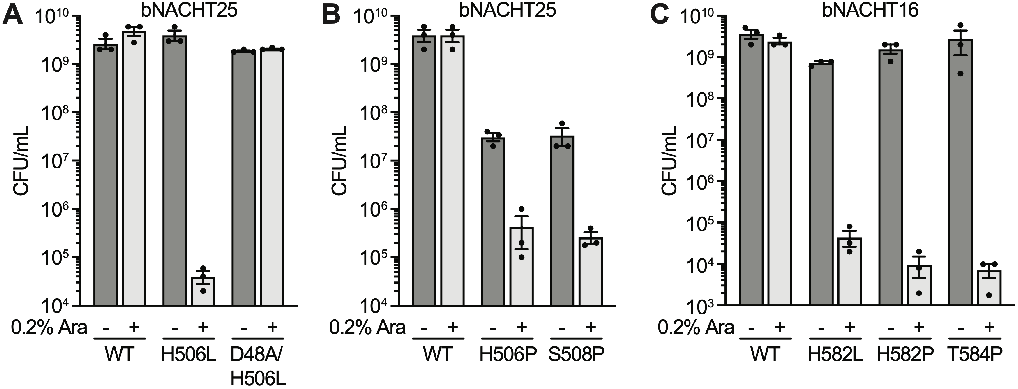
Human disease-associated mutations hyperactivate bacterial NACHT proteins. **(A)–(B)** Quantification of colony formation of *E. coli* expressing wild-type (WT) bNACHT25 or alleles with the indicated mutations. **(C)** Quantification of colony formation of *E. coli* expressing bNACHT16 with the indicated mutations. See **Figure S9** for an alignment of NLRC4, bNACHT16, and bNACHT25. For A–C, gene expression was induced with arabinose. Symbols denote induction (+) or lack of induction (-). Data represent the mean ± s.e.m. of *n* = *3* biological replicates, shown as individual points.

The disease-associated mutation at S445 also results in hyperactivation of NLRC4 (Romberg et al., 2017; Volker-Touw et al., 2017). To test whether we could recapitulate the effects of mutation to this residue in bacteria, we mutated the corresponding residue in bNACHT25 (S508P) and bNACHT16 (T584P). Overexpression of both the bNACHT25 and the bNACHT16 mutants resulted in inhibition of bacterial growth (**Figure 5B and C**). These data demonstrate that NACHT modules in humans and bacteria can be hyperactivated by similar mutations, suggesting these proteins have a similar mechanism of effector domain activation.

Our analysis of *orf008* and *orf015* from phage T5 demonstrated that these proteins alter bNACHT signaling (**Figure 4**). To further characterize the effect of these two phage genes on bNACHT activity, we co-expressed *orf008* and *orf015* with the hyperactive allele of bNACHT25, then measured bacterial growth. As predicted from our previous analysis, *orf008* did not appreciably alter colony formation as hyperactive bNACHT25 already leads to bNACHT-dependent growth inhibition (**Figure 5A**, **Figure 6**). However, the *orf015* gene was sufficient to rescue the growth inhibition of the bNACHT25 hyperactive allele (**Figure 6A**). These data are consistent with the impact of *orf015* on bNACHT25 signaling during infection (**Figure 4C**). Similar results were also obtained for bNACHT01 when this protein was hyperactivated by overexpression, a method used to activate other antiphage systems (**Figure 6C**) (Severin et al., 2018). These data demonstrate that *orf015* interrupts bacterial NLR signaling even in the absence of phage.

**Figure 6.**
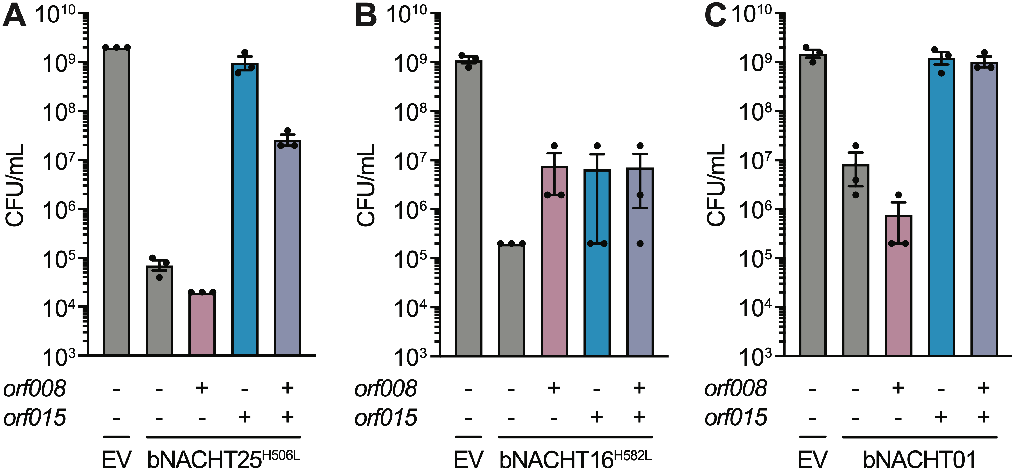
Phage proteins alter the toxicity of hyperactive bacterial NACHT proteins. **(A)–(C)** Quantification of colony formation of *E. coli* expressing a bNACHT gene with the indicated genotype on one plasmid and the indicated phage T5 gene(s) on a second plasmid. Expression of *orf008, orf015*, and *mCherry* was induced with IPTG. Symbols denote induction of an *mCherry* negative control gene (-) or induction of the indicated phage gene (+). Expression of the indicated bNACHT gene or empty vector (EV) was arabinose-inducible. Data represent the mean ± s.e.m. of *n* = *3* biological replicates, shown as individual points.

## Discussion

Here we identify that NACHT module-containing proteins are abundant and widespread in the genomes of bacteria where they are potent phage defense systems. Bacterial and animal NACHT proteins are highly similar in their overall domain architecture, the predicted structure of their NACHT module, and their role in immune signaling. These data establish bacterial NACHT proteins are related to eukaryotic NLRs. In support of a shared molecular mechanism of NACHT module activation, point mutations that hyperactivate NACHT modules in human cells also hyperactive NACHT modules in bacteria. Hyperactivated alleles of bacterial NACHT proteins inhibited growth of bacteria. Further, phage infection also appeared to inhibit growth of NACHT protein-expressing bacteria, suggesting that these systems may inhibit phage replication via abortive infection. Abortive infection is a form of programmed cell death that interrupts the viral lifecycle by prematurely destroying a host component essential to virion production. In this way, the antimicrobial signaling outcome of bacterial NLR-related proteins may also be similar to mammalian inflammasomes, which initiate a caspase-dependent programmed cell death called pyroptosis when activated. We anticipate that further understanding of the molecular mechanisms of bacterial NACHT protein signaling will provide valuable insights into human NLRs.

Our expansive bioinformatic analysis found that bacteria encode the largest diversity of NACHT module sequences compared to other superkingdoms, which suggests that this protein module first evolved in bacteria before being acquired into the genomes of eukaryotes. However, not all eukaryotic NACHT module sequences are monophyletic and each often clusters with distinct groups of bacterial NACHT proteins, implying that horizontal gene transfer of NACHT modules from prokaryotes to eukaryotes has occurred on multiple occasions. Evidence for one transfer event is found in NACHT module Clade 12, which groups mammalian NLRs (aka Caterpillar genes) with bacterial NACHT proteins from *Rickettsiales*, an order of intracellular bacteria. This observation suggests that metazoans acquired their NLRs from *Rickettsiales*. A similar horizontal gene transfer event has been suggested for the innate immune gene STING, however the most probable bacterial source for that event is the Bacteroidetes (Burroughs and Aravind, 2020). Both Bacteroidetes (living extracellularly as a symbiont) and *Rickettsia* (living intracellularly) have intimate interactions with eukaryotes yet distinctly different lifestyles. The shared evolutionary history of NACHT genes may enable future investigators to take advantage of studying bacterium-phage interactions to learn about cryptic aspects of human NLR signaling.

Fungi also encode NACHT proteins that are uniquely suited to their lifestyle. The HET-D and HET-E proteins from the filamentous fungus *Podospora anserina* are NACHT proteins that mediate kin recognition after two cells have fused their cytoplasms. When kin cells expressing these proteins fuse, the subsequent heterokaryon survives, however, when non-kin cells expressing HET-D or HET-E fuse, the NACHT protein initiates programmed cell death (Aanen et al., 2010; Glass and Dementhon, 2006). HET-D/E recognize allelic differences in the HET-C protein to distinguish kin, i.e., self from non-self (Espagne et al., 2002). This phenomenon is known as heterokaryon incompatibility. In related systems, heterokaryon incompatibility has been shown to restrict the spread of endogenous viruses between non-kin fungi (Choi et al., 2012; Paoletti and Saupe, 2009). Thus, in fungi, as in animals and bacteria, NLR-related proteins are part of the innate immune system.

NLRs within the mammalian inflammasome require additional factors to induce cell death. NLRC4 requires the pore-forming protein Gasdermin D to execute cell death (pyroptosis) (Broz and Dixit, 2016; Shi et al., 2015). Gasdermin D homologs can also be found in fungi, where they mediate heterokaryon incompatibility, and in bacteria, where they mediate antiphage signaling (Clavé et al., 2022; Daskalov et al., 2020; Johnson et al., 2022). However, bacterial NLR-related proteins do not require Gasdermin D homologs for signaling, unlike the mammalian NLRs which activate cell death via Gasdermin D. Heterokaryon incompatibility loci are highly polymorphic across fungi, and there are many more than *het-d/e* (NLRs) and *rcd-1* (Gasdermin D) (Dyrka et al., 2014; Van der Nest et al., 2014). These observations suggest that heterokaryon incompatibility loci, like bacterial antiphage systems, may be an important repository for identifying mammalian innate immune genes.

Bacterial NACHT proteins are the first example of an innate immune antiphage system in bacteria capable of defending against RNA phages. While adaptive immune systems like CRISPR can be programmed to defeat RNA phages (Strutt et al., 2018), this may not represent their natural function. Bacterial NLRs capable of recognizing RNA phages also recognize DNA phages, suggesting that the stimulus recognized is highly conserved between disparate viruses. We do not yet know what the stimulus might be, or if the stimulus is the same for all bacterial NACHT proteins. However, we are able to synthetically activate these proteins using mutations that hyper-activate mammalian NLRs. Many NACHT-associated effector domains are highly conserved and found across multiple known and predicted antiphage systems but remain as yet biochemically uncharacterized. Given that they cannot be readily activated in the absence of a phage (which might be unknown), synthetic activation might prove highly useful to study the large array of effector domains fused to the N-terminus of bacterial NACHTs. Some noteworthy examples include: (i) the Schlafen RNase domain found at the N-terminus of bNACHT34 that is related to human Schlafen proteins involved in HIV1 restriction (Jakobsen et al., 2013); (ii) The PNPase domain that is predicted to generate NAD^+^ or other nucleotide derivatives (Burroughs and Aravind; Burroughs et al., 2015); (iii) bacterial domains related to the Death-superfamily domains found in metazoan apoptosis (Kaur et al., 2020, 2021).

Our data support a unifying role for proteins encoding NACHT modules and related STAND NTPases as mediators of innate immunity across the tree of life. NACHT module-encoding NLRs in mammals initiate inflammation and are potently antimicrobial. Fungal NACHT proteins mediate heterokaryon incompatibility, which can stop viral transmission. Here, we demonstrate that the bacterial NACHT proteins are antiphage. Land plants also show an expansion of the antibacterial and antiviral R (NB-ARC) proteins that contain another clade of STAND NTPase modules, i.e., the AP-ATPase, which is a sister-group of the NACHT clade. Thus, it appears that the NACHT and related STAND modules define a biochemical scaffold that is especially suited for immune signaling. One potential explanation is that these modules can serve as switches that combine pathogen sensing, activation-threshold setting, signal transduction, and effector deployment, all in a single protein (Lisa et al., 2019; Marquenet and Richet, 2007). Understanding the unique qualities of the NACHT module is an exciting area for future investigation.

## Supplementary Discussion

In this discussion, we suggest possible functions and evolutionary implications of the diverse array of domain architectures identified in bacterial NACHT proteins. First, we expand upon the types of sensing domains found in bacterial NACHT proteins. These sensor domains tend to be located C-terminally of the NACHT module.

1. *“Antigen receptor type”:* These are typified by a C-terminal domain that is likely to bind a host protein, thereby “guarding” an essential host process (Biezen and Jones, 1998; van Wersch et al., 2020), or a protein/other molecule from the phage, thereby directly recognizing infection. These can be further divided into 2 types:

i. The FGS domain type – this has a single FGS domain that in several cases likely undergoes directed diversification by the DGR Reverse transcriptase (Kaur et al., 2020, 2021). Thus, they may be potential bacterial analogs of metazoan adaptive antigen-receptors like antibodies or TCRs.
ii. Those with superstructure forming repeats, namely LRRs, TPRs, HEATs, β-propellers, or pentapeptide repeats (Tørresen et al., 2019).These are often predicted to form toroid or helical structures that may recognize phage components, comparable to pathogen associated molecular pattern (PAMP) recognition in the metazoan NLRs that form the inflammasome (Li and Wu, 2021). Indeed, eukaryotic NACHTs appear to have been derived on multiple occasions from bacterial versions with this type of sensor domain.
2. *“Short C-terminal extensions”:* In this group, the C-terminus has a shorter domain rather than an FGS or superstructure forming element. bNACHT01 and its homologs from diverse bacteria form one of the largest monophyletic clades of bacterial NACHT proteins with this type of sensor domain.
3. *“Hybrids”:* These have a C-terminal superstructure forming module in addition to a small distinct C-terminal domain that may tether the bacterial NACHT protein to some endogenous component.
4. *“TM regions”:* These are typified by multiple TM regions (sometimes inserted into the NACHT module). A subset of bacterial NACHT proteins with this type of sensing domain likely recognize intra-membrane or periplasmic signals (Kaur et al., 2020, 2021).
5. *“Other C-terminal sensor domains”:* This group is characterized by a C-terminal domain that is distinct from those described above with a predicted sensory role. In cyanobacteria, a common example of this type has a GUN4 domain that senses chlorophyll derivatives and is predicted to monitor photosynthesis status or virally induced changes to photosynthesis (Kaur et al., 2020, 2021; Larkin et al., 2003).

With the exception of the bNACHT01-like clade (Clade 14, **Figure 2**), bacterial NACHT proteins are characterized by fusions to a remarkable array of effector domains that most frequently occur at the extreme N-terminus before the NACHT module and more infrequently at the extreme C-terminus after the repeat or FGS domains. The effectors found on bacterial NACHT proteins can be categorized into two types: enzymatic effectors and adaptors.

1. *“Enzymatic effectors”:* These are effectors that are predicted to have a catalytic role within the cell that may directly mediate cell death or inhibition of phage replication through depletion of an essential resource.

i. RNases (e.g., Schlafen (Burroughs and Aravind, 2020) and HEPN metal-independent RNase domains (Anantharaman et al., 2013))
ii. DNases (e.g., REase (restriction endonuclease fold) and HNH domains)
iii. NAD^+^-targeting domains (e.g., TIR, Sirtuin (SIR2), DrHyd, PNPase) (Burroughs and Aravind; Burroughs et al., 2015)
iv. Nucleotide-signal-generating/degrading enzymes (e.g., cNMP-cyclases, alarmone synthetases, SMODS, calcineurin-like phosphoesterases, a novel pol-β fold nucleotidyltransferase) (Burroughs and Aravind; Burroughs et al., 2015)
v. Peptidases (e.g., Caspase, Trypsin, transglutaminase-like papain-like peptidase and the deSUMOylating peptidases like SMT4) (Kaur et al., 2020, 2021; Nayak and Müller, 2014).
vi. Other potential protein-modifying enzymes (e.g., protein (S/T/Y) kinase, GT4 family glycosyltransferases) (Kaur et al., 2020, 2021).
2. *“Adaptors”:* These effectors without clear catalytic functions that may inhibit phage replication by recruiting other, endogenous host effectors or activating a transcriptional response.

i. Effector Associated Domains and Death-like domains (Kaur et al., 2020, 2021)
ii. Transcription regulatory/DNA-binding domains (e.g., wHTH domains)
iii. RNA-binding domains (e.g., S1/Coldshock OB-fold domain)
iv. Cyclic nucleotide sensors (e.g., cNMPBD)
v. Chlorophyll-precursor sensors (e.g., GUN4 (Larkin et al., 2003))

While most of the effector domains on bacterial NACHTs are found at the N-terminus of the protein, before the core NACHT module, some occur at the extreme C-terminus of the protein, after tandem repeat or FGS domains. Structural considerations based on published structures of metazoan NLRs suggest that, in these cases, the C-terminal effectors are likely in a similar position as the N-terminal effectors in the folded polypeptide. We predict that the toroidal or helical elements of the C-terminus may allow the effector domain to maintain its normal spatial location, despite being located at the opposite terminus of the protein.

Overall, the numbers of NACHT domains encoded by an organism shows a power law distribution that might be fitted by y=x^-2.05^, with the largest number of paralogs found in cyanobacteria (up to 23 per genome; **Figure S2**). Phylogenetic analysis suggests that proteins with more closely related NACHT modules tend to have similar C-terminal sensor domains. The effector domains to which the NACHT module is fused can vary dramatically between species of the same genus or bacterial lineage, suggesting that the effector domains are in an arms race with viral inhibitors that target them.

## Supporting information

Kibby et al. Table S1

Kibby et al. Table S2

Kibby et al. Table S3

Kibby et al. Table S4

Kibby et al. Table S5

Kibby et al. Table S6

Kibby et al. Table S7

## Acknowledgements

The authors would like to thank R. Vance for advice; A. Litt for assistance with cloning; the CU Boulder Department of Biochemistry Shared Instruments Pool core facility (RRID:SCR_018986) and its staff; A. Scott and K.E. Jackson at the University of Colorado Boulder BioFrontiers Institute’s Next Generation Sequencing Core (RRID:SCR_019308) for technical assistance; Brian Kvitko for generously sharing strains and plasmids; and members of the Whiteley lab for advice and helpful discussion. This work was funded by a Mallinckrodt Foundation Grant (A.T.W.). E.M.K is supported in part by the NIH T32 Signaling and Cellular Regulation training grant (T32 GM008759 and T32 GM142607). AMB and LA are supported by the Intramural Research Program of the National Library of Medicine at the NIH.

## Author contributions

Experiments were designed and conceived by E.M.K., A.N.C., and A.T.W. Bioinformatic identification and characterization of bacterial NACHTs was performed by L.A. and A.M.B. E.M.K. and A.T.W. selected bNACHT candidates for further analysis. Phage spotting assays were performed by E.M.K. and A.N.C. Phage suppressor generation and sequencing was performed by E.M.K. T5 ORF coexpression assays were performed by E.M.K. and T.A.N. Hyperactive bNACHTs were designed and tested by A.N.C. Figures were prepared by E.M.K., J.A.V, and A.M.B. The manuscript was written by E.M.K. and A.T.W. All authors contributed to editing the manuscript and support the conclusions.

## Declaration of interests

The authors declare no competing interests.

Correspondence and requests for materials should be addressed to the lead contact, Aaron Whiteley

### Materials availability

Strains, plasmids, and phages used in this study are available upon reasonable request.

## Materials and Methods

### Bacterial strains and culture conditions

The *E. coli* used in this study are listed in **Table S6**. *E. coli* were cultured in LB medium (1% tryptone, 0.5% yeast extract, and 0.5% NaCl) shaking at 37 °C and 220 rpm in 1-3 mL of media in 14 mL culture tubes, unless otherwise indicated. Where applicable, carbenicillin (100 μg/mL), chloramphenicol (20 μg/mL), and tetracycline (15 μg/mL) were added. We defined “overnight” bacterial cultures as 16-20 hours post-inoculation from a glycerol stock or single colony. All strains were frozen for storage in LB plus 30% glycerol at −70 °C. *E. coli* OmniPir was used for construction and propagation of all plasmids. *E. coli* MG1655 (CGSC6300) was used to collect all experimental data.

*E. coli* OmniPir was constructed from OmniMAX 2 T1^R^ *E. coli* (Thermo Fisher Scientific) and *pGRG36pir-116* as previously described (Kvitko et al., 2012). Briefly, the *pir116* gene was integrated at the Tn7 attachment site by conjugating *pGRG36pir-116* into OmniMAX *E. coli*, cultivating bacteria at the permissive temperature with arabinose induction, then curing the plasmid at 42 °C. Integration of *pir116* was confirmed by PCR and retention of the F’ plasmid was confirmed by tetracycline resistance. *E. coli* MG1655 F+ strain was constructed by isolating the F’ plasmid from OmniPir following a previously described protocol (Herrick et al., 2018) and electroporating the purified plasmid into electrocompetent MG1655, followed by selection with tetracycline.

MMCG medium (47.8 mM Na_2_HPO_4_, 22 mM KH_2_PO_4_, 18.7 mM NH4Cl, 8.6 mM NaCl, 22.2 mM Glucose, 2 mM MgSO_4_, 100 μM CaCl_2_, 3 μM Thiamine, Trace Metals at 0.1× (Trace Metals Mixture T1001, Teknova, final concentration: 5mM Ferric chloride, 2mM Calcium chloride, 1mM Manganese chloride, 1mM Zinc Sulfate, 0.2mM Cobalt chloride, 0.2mM Cupric chloride, 0.2mM Nickel chloride, 0.2mM Sodium molybdate, 0.2mM Sodium selenite, 0.2mM Boric acid)) with appropriate antibiotics was used to collect all experimental data. When experiments required bacteria expressing two plasmids, strains were grown using reduced antibiotic concentrations to enhance growth rate (MMCG with 20 μg/mL carbenicillin and 4 μg/mL chloramphenicol).

When growing strains that required induction, 100 μM IPTG or 0.2% arabinose was used to induce, as appropriate.

### Phage amplification and storage

The phages used in this study are listed in **Table S7**. Phages were amplified via either liquid or plate amplification using a modified double agar overlay (Kropinski et al., 2009). For liquid amplification, 5 mL mid-log cultures of *E. coli* MG1655 in LB plus 10 mM MgCl_2_, 10 mM CaCl_2_, and 100 μM MnCl_2_ were infected with phage at an MOI of 0.1 and grown, shaking, for 2-16 hours. The supernatant was harvested and filtered through a 0.2 μm spin filter to remove bacterial contamination. For plate amplification, 400 μL of mid-log MG1655 were mixed with 3.5 mL LB soft agar mix (LB with 0.35% agar and 10 mM MgCl_2_, 10 mM CaCl_2_ and 100 μM MnCl_2_) and 100-1,000 PFU. Plates were then incubated for 16 hours at 37 °C. 5 mL of SM buffer (100 mM NaCl, 8 mM MgSO_4_, 50 mM Tris-HCl pH 7.5, 0.01% gelatin) was added to the plate and allowed to soak out the phages for 1 hour before SM buffer was collected and passed through a 0.2 μm filter. All phages were stored at 4 °C in SM buffer or LB.

### Plasmid construction

The plasmids used in this study are listed in **Table S6**. DNA manipulations and cloning were performed as previously described (Whiteley et al., 2019). Briefly, genes of interest were amplified from phage or bacterial genomic DNA using Q5 Hot Start High Fidelity Master Mix (NEB, M0494L), or synthesized as GeneFragments (Genewiz) flanked by ≥ 18 base pairs of homology to the vector backbone. Ligation of genes into restriction-digested, linearized vectors was accomplished using modified Gibson Assembly (Gibson et al., 2009). Gibson reactions were transformed via heat shock or electroporation into competent OmniPir and plated onto appropriate antibiotic selection. Where possible, bNACHT coding sequences and endogenous regulatory regions were amplified from the genomic DNA of *E. coli* strains from the ECOR collection (Ochman and Selander, 1984). All other bNACHT gene inserts were ordered as GeneFragments (Genewiz). bNACHT point mutations were generated by amplifying out the gene of interest in two parts from a plasmid template, with the desired mutation occurring in the overlapping region between the two amplicons. Inserts for expression of all *orf008* and *orf015* alleles were amplified from appropriate phage genomic DNA. Unless otherwise indicated, all enzymes were purchased from New England Biolabs.

For all vectors using the pLOCO2 backbone, pAW1382 was amplified and purified from OmniPir. Purified plasmid was then linearized using SbfI-HF and NotI-HF or FseI-HF. Gibson ligation was used to circularize the plasmid with a new insert.

For all vectors using the pTACxc backbone, pAW1608 was amplified and purified from OmniPir. Purified plasmid was then linearized using BamHI-HF and NotI-HF. Gibson ligation was used to circularize the plasmid with the new insert.

For all vectors using the pBAD30x backbone, pAW1367 was amplified and purified from OmniPir. Purified plasmid was then linearized using EcoRI-HF and HindIII-HF. Gibson ligation was used to circularize the plasmid with the new insert.

Sanger sequencing (Genewiz) was used to validate the correct sequence within the multiple cloning site. Additionally, all plasmids expressing novel bNACHT genes were sequence verified by Illumina sequencing (CU Boulder Sequencing Facility). A NextSeq V2 Mid Output 150-cycle kit was used to sequence the plasmids. Reads were mapped to the predicted plasmid sequence using the *Map to Reference* feature of Geneious Prime (default settings).

### Efficiency of plating/phage resistance analysis

A modified double agar overlay was used to measure the efficiency of plating (EOP) of phages (Kropinski et al., 2009; Ledvina et al., 2022). Briefly, overnight cultures of *E. coli* MG1655 expressing the indicated plasmids cultured in MMCG plus appropriate antibiotics were diluted 1:10 into the same media and cultivated for an additional two hours to reach mid-log phase (OD_600_ 0.1-0.8). 400 μL of the mid-log culture was mixed with 3.5 mL MMCG (0.35% agar), plus an additional 5 mM MgCl_2_ and 100 μM MnCl_2_. The mixture was poured onto an MMCG (1.6% agar) plate and cooled for ~15 minutes. 2 μL of a phage dilution series in SM buffer was spotted onto the overlay and allowed to adsorb for 10 minutes before incubating the plate overnight at 37 °C.

Plaque formation was enumerated the following day. For instances with a hazy zone of clearance rather than individual plaque formation, the lowest phage concentration at which clearance was observed was counted as ten plaques. In instances where no clearance or plaque formation was visible, 0.9 plaques at the least dilute spot were used as the limit of detection.

Fold protection was calculated using the inverse of EOP. The PFU of a given phage lysate was measured on sensitive host bacteria, expressing an empty vector, then divided by the PFU for the same phage lysate measure on test bacterial strains. In this way, a 10-fold decrease in EOP is a 10-fold increase phage protection.

bNACHT22 is included in **Figure S4** but not selected for inclusion in **Figure 3**. Although we did observe a decrease in T3 PFU for this system, we did not observe an expected decrease in T3 plaque size, which undermined our confidence in the significance of this result.

### Time course of phage infection

Overnight cultures of the appropriate strains were inoculated in 30 mL MMCG plus 100 μg/mL carbenicillin, 5 mM MgCl_2_, and 100 μM MnCl_2_ to an OD_600_ of 0.1. Cultures were then cultivated shaking at 37°C for two hours and infected with phage at the indicated MOI. Culture OD_600_ was measured at indicated times.

### Identification of bacterial NACHTs

We started with an initial sequence library of known NACHT modules from prior studies (Aravind et al., 2004; Kaur et al., 2021; Leipe et al., 2004). Upon identification of new homologs these were then integrated into the initial library for further large-scale sequence analysis as described below. We iterated this procedure for several rounds, and eventually generated an exhaustive collection of NACHT modules homologs. To detect distant relationships, iterative sequence profile searches were conducted using the PSI-BLAST (RRID:SCR_001010) (Altschul et al., 1997) and JACKHMMER (RRID:SCR_005305) (Potter et al., 2018) programs with profile-inclusion threshold of expect (e)-value at 0.005 against the non-redundant database of National Center for Biotechnology Information (NCBI) clustered down to 50%. Clustering of proteins based on bit score density and length of aligned sequence was performed using the BLASTCLUST program (ftp://ftp.ncbi.nih.gov/blast/documents/blastclust.html). Remote homology searches were performed using profile-profile comparisons with HHpred program (RRID:SCR_010276) (Zimmermann et al., 2018) against profile libraries comprised of the PFAM (RRID: SCR_004726) (Mistry et al., 2020) and PDB (RRID:SCR_012820) (Berman et al., 2000) databases as well as an in-house library of profiles of conserved domains. Multiple sequence alignments were built using the Kalign (RRID:SCR_011810) (Lassmann, 2019) and Muscle (RRID:SCR_011812) (Edgar, 2004) programs followed by manual adjustments based on profile–profile alignment, secondary structure prediction, and structural alignment.

Searches for establishing taxonomic counts of NACHT domains from lineages across the tree of life and viruses was performed using a custom database of 14785 completely sequenced genomes (6847 bacteria) using known NACHT domains as queries for PSI-BLAST searches run for 3 iterations with an inclusion threshold of 0.0001. The detected candidates were then run through a confirmation step with the RPS-BLAST program to obtain the final count of NACHT proteins.

Secondary structures were predicted using the JPred (RRID:SCR_016504) (Drozdetskiy et al., 2015) program. Phylogenetic analysis was conducted using the maximum likelihood method implemented in the IQtree program (RRID: SCR_017254) (Minh et al., 2020) under multiple parameter regimes using: 1) the Q.pfam substitution matrix derived from alignments in the Pfam database and 1 invariant site category with 8 gamma distributed sites; 2) the LG substitution matrix with 1 invariant site category with 8 gamma distributed sites or 3) with a 20-profile mixture model. The trees were rendered using the FigTree program (RRID:SCR_008515) (http://tree.bio.ed.ac.uk/software/figtree/).

### Domain detection

To establish the domain architectures of the NACHT proteins, they were first searched for previously known domains using the RPS-BLAST program with the Pfam database and a custom database including all of domains detected by the Aravind group and augmentations of the Pfam profiles to improve detection. Unknown regions were then investigated. Profile-profile searches were performed with the HHpred program against libraries of profiles based on non-redundant PDB structures, the Pfam database, and a custom collection of profiles of domains not detected by Pfam. Kalign with default parameters and Mafft with maxiterate= 3000, globalpair, op= 1.9 and ep= 0.5 were used to generate input multiple sequence alignments (MSA), followed by refinements using HHpred profile-profile matches or HMM-align. For specific cases structural modeling was performed using the RoseTTAFold program, which uses a “three-track” neural network, utilizing patterns of sequence conservation, distance inferred from coevolutionary changes in MSAs, and coordinate information (Baek et al., 2021). MSAs of related sequences (>30% similarity) were used to initiate HHpred searches for the initial step of correlated position and contact identification to be used by the neural networks.

### Analysis of differential diversity of the NACHT module and the SNaCT domain

The analysis of the Shannon entropy (H) for a given multiple sequence alignment was performed using the equation:

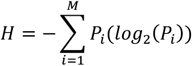

P is the fraction of residues of amino acid type i and M is the number of amino acid types. The Shannon entropy for the ith position in the alignment is ranges from 0 (only one residue at that position) to 4.32 (all 20 residues equally represented at that position).

Trident entropy was used as the metric used to analyze the differential divergence of the NACHT module and the SNaCT domain in clade 14. This measure simultaneously unite three distinct elements (hence trident) of positional variability (Valdar, 2002) namely: 1. residue diversity; 2. Biochemical diversity among residues; 3. Gapiness of an alignment column. The first t(x) is measured using normalized Shannon entropy (see above); the second r(x) is measured using dissimilarity between two amino acids based on Karlin’s formula using a substitution matrix computed from the alignment; the third g(x) measures the number of gaps in the column. The three united as a product (S=t(x)^a^.r(x)^b^.g(x)^c^, with each factor scaled with an exponent. The respective exponents used here are: a=1, b=½ and c=3. The analysis of the entropy values which were thus derived were performed in the R language.

### bNACHT library preparation

bNACHT proteins were selected for screening by considering relatedness of the source genome to *E. coli* and protein domain diversity. For each gene tested, we included the coding sequence of the bNACHT gene, as well as any other genes in the operon. We also included the endogenous regulatory elements of each system, using bPROM (Solovyev and Salamov, 2011) to predict bacterial promoters and ARNOLD to predict terminators (Naville et al.). We included at least 100 nucleotides to the 3’ and 5’ region of the gene of interest, to ensure that even unidentified regulatory elements would be included.

### Growth inhibition measurements

The impact of bNACHT expression with and without *orf008* and *orf015* alleles on bacterial growth was quantified using a colony formation assay. *E. coli* was cultivated overnight in MMCG with appropriate antibiotics. Cultures were diluted in a 10-fold series into MMCG and 5 μL of each dilution was spotted onto a MMCG agar plate containing the appropriate antibiotics, as well as IPTG and/or arabinose as appropriate. Spotted bacteria were allowed to dry for ~10 minutes before the plates were incubated overnight at 37 °C.

Growth inihibition was measured the following day by enumerating the colony forming units of each strain, reported as CFU/mL for the starting culture. For instances where bacteria were growing but no individual colonies could be counted, the lowest bacterial concentration at which growth was observed was counted as ten CFU. In instances where no growth was visible, 0.9 CFU at the least dilute spot was used as the limit of detection.

### Phage suppressor generation and amplification

T5 phages able to evade bNACHT01-medaited protection were generated by mixing 400 μL of mid-log bacteria expressing bNACHT01 in MMCG plus 100 μg/mL carbenicillin with wild-type T5 at an MOI ~10 and pouring the mixture onto a MMCG agar plate. Individual plaques were isolated and spot-plated onto *E. coli* MG1655 expressing bNACHT01 to confirm that phages were able to replicate in the presence of bNACHT01 and to plaque-purify each clone. Phage bNACHT01 suppressors were generated using three separate wild-type T5 stocks amplified from individual plaque purifications. Phage T5 suppressors were subsequently plate amplified on *E. coli* MG1655 expressing bNACHT01 in MMCG.

### Genome sequencing and analysis of phage suppressors

Phage genomes were sequenced as previously described (Millman et al., 2020). Briefly, 450 μL of phage lysate (>10^7^ PFU/mL) was treated with DNAse I (final concentration 3 × 10^-3^ U/μL) and RNAse A (final concentration 3 × 10^-2^ μg/μL) and incubated for 1.5 hours at 37 °C to remove extracellular nucleic acids. EDTA was added (final concentration 20 mM) to stop the reaction. Phage genomes were subsequently isolated and purified using the Qiagen DNeasy cleanup kit, starting at the proteinase K digestion step (Millman et al., 2020). Extracted phage genomes were prepared for Illumina sequencing using a modification of the Nextera kit protocol as previously described (Baym et al., 2015). Illumina sequencing was performed using a MiSeq V2 Micro 300-cycle kit (CU Boulder Sequencing Facility). Reads were mapped to Genome accession AY587007 (empirically determined to be most similar to the T5 phage used in this study) using Geneious software’s *Map to Reference* feature. Reads were trimmed to remove the Nextera adapter sequences before mapping (sequence trimmed: AGATGTGTATAAGAGACAG) using the “Trim primers” option, with otherwise default settings. Sequences were mapped using default settings, selecting “map multiple best matches to all locations” to accommodate repetitive T5 sequences.

Geneious was also used for variant detection from the reference T5 genome. Variants that were present in ≥75 percent of reads from the suppressor phage genome but not the parent phage genome were identified as potential suppressor mutations.

### Effect of phage genes on bNACHT protection against phage

Bacterial strains were cultured overnight in MMCG plus 20 μg/mL carbenicillin and 4 μg/mL chloramphenicol. Cultures were then diluted 1:10 into the same media with or without 100 μM IPTG and grown for 4 more hours to reach midlog phase. Phage resistance was measured as described above, with the addition of IPTG to the MMCG top agar (0.35%) to continue inducing conditions.

### Data visualization and statistics

All experiments were performed in biological triplicate using cultures grown on three separate days. Data was plotted using Graphpad Prism 9 at an n of 3 with error bars indicating standard error of the mean. Illumina sequencing results were analyzed using Geneious Prime Software. Geneious Prime was also used to generate alignments, using MAFFT alignment (Katoh et al., 2002) and default settings. Figures were created using Adobe Illustrator CC.

## Data availability

The data supporting the findings of this study are available within the article and its supplementary files.

## Supplementary Data Tables

Table S1-Diverse NACHT proteins bioinformatically identified

Table S2-Experimentally investigated bacterial NACHT proteins

Table S3-Rate of suppressor isolation for T5 infecting bNACHT01

Table S4-T5 suppressor mutations in *orf008* and *orf015*

Table S5-All T5 suppressor mutations

Table S6-*E. coli* strains and plasmids used in this study

Table S7-Bacteriophages used in this study

## Supplementary Figures

**Figure S1.**
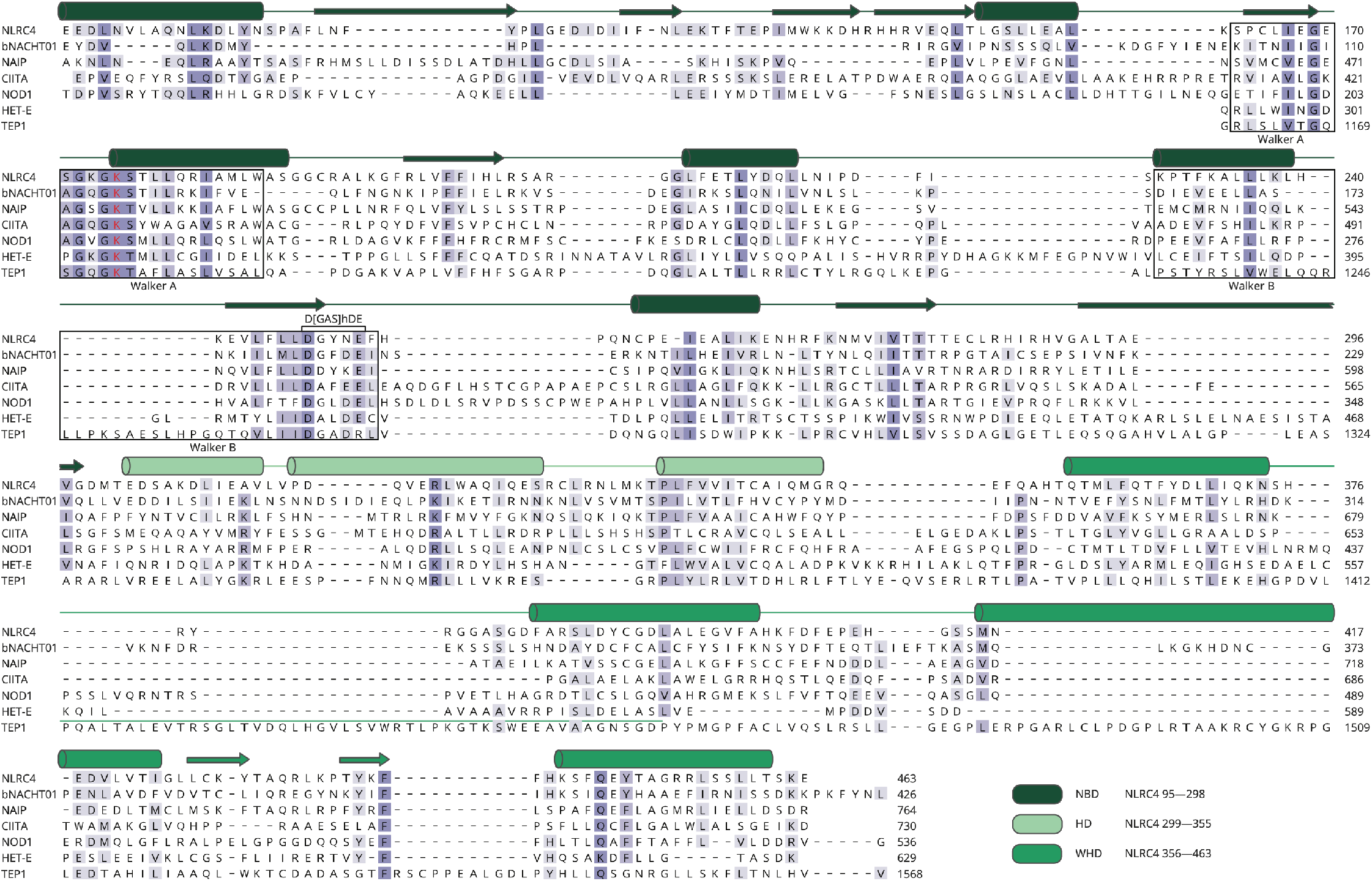
Protein alignment of the NACHT modules from bNACHT01 and other NACHT proteins. Protein alignment of the NACHT modules of NLRC4 (*Mus musculus;* NP_001028539), bNACHT01 (*Klebsiella pneumoniae* MGH 35; WP_015632533), NAIP (*Homo sapiens;* A55478), CIITA (*Homo sapiens;* AAA88861), NOD1 (*Homo sapiens;* AAD29125), HET-E (*Podospora anserina;* Q00808), and TEP1 (*Homo sapiens;* AAC51107). The secondary structure of NLRC4 as determined using structure 4KXF in the PDB (Hu et al., 2013), is indicated above with alpha helices depicted as cylinders and beta sheets depicted as arrows. Secondary structure elements are color coded to represent the NBD, HD, and WHD of NLRC4. Amino acid residues are color-coded based on conservation such that darker colors indicate a higher degree of conservation in this alignment. Black boxes indicate the Walker A and Walker B motifs. The conserved lysine in the Walker A motif is highlighted in red. The D[GAS]hDE motif within the Walker B region that distinguishes NACHT modules from other STAND NTPases is indicated.

**Figure S2.**
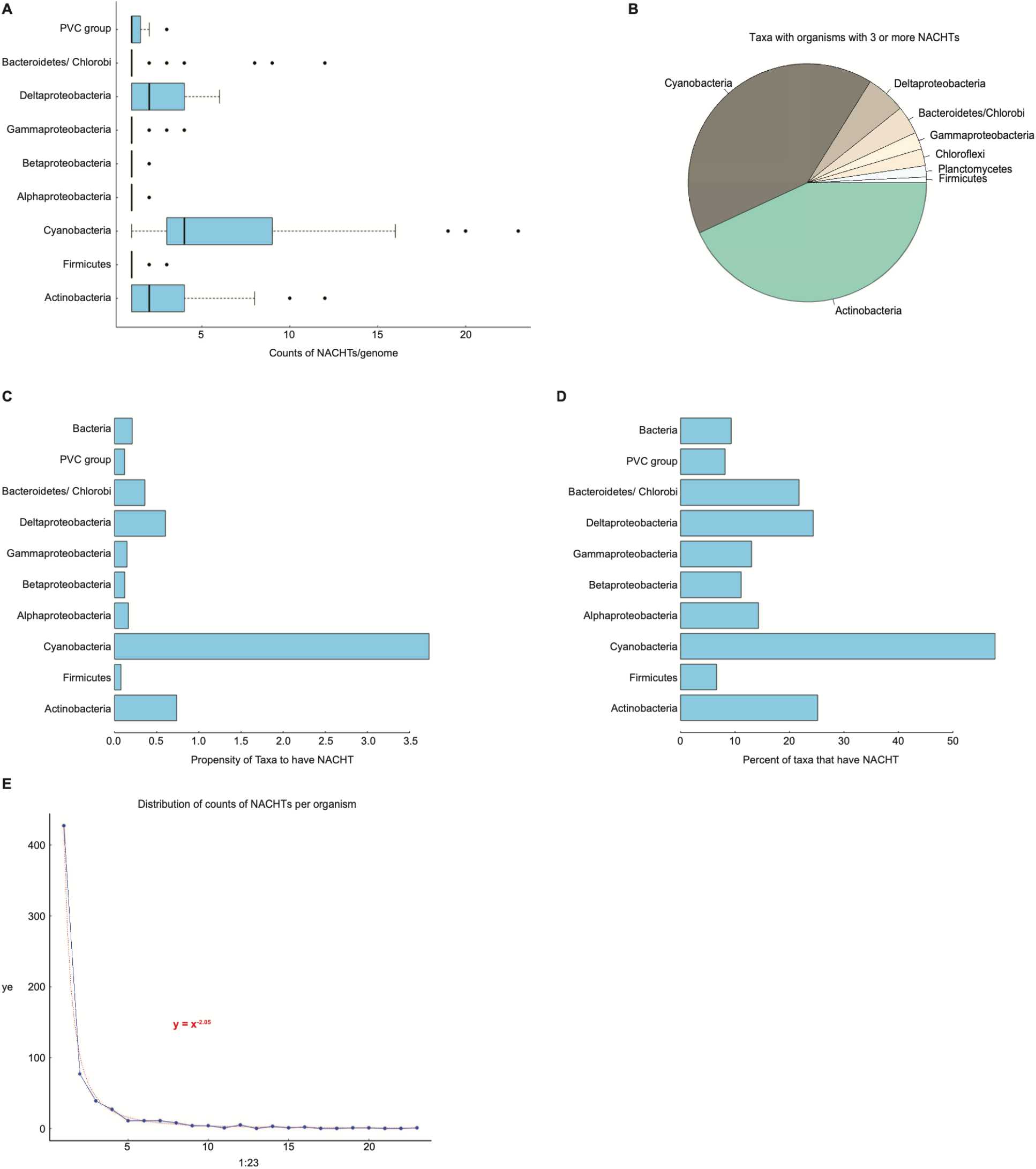
Distribution of NACHT proteins in bacterial taxa. **(A)** Quantification of the number of individual NACHTs found within a single genome across different bacterial taxa. The maximum for the x-axis is 23. Plantomycetota, Verrumicrobiota, Chlamydiota (PVC). **(B)** Relative distribution of taxa that have organisms with 3 or more NACHTs in a single genome. **(C)** Propensity of organisms within the indicated taxa to encode a NACHT module-containing protein. **(D)** Percent of organisms in the indicated taxa to encode at least one NACHT module-containing protein. **(E)** Distribution of the number of NACHT proteins per organism, which can be fitted to the equation y = x^-2,05^. For A–E, a custom database of complete bacterial genomes was used to analyze distribution of NACHT proteins in diverse bacterial taxa.

**Figure S3.**
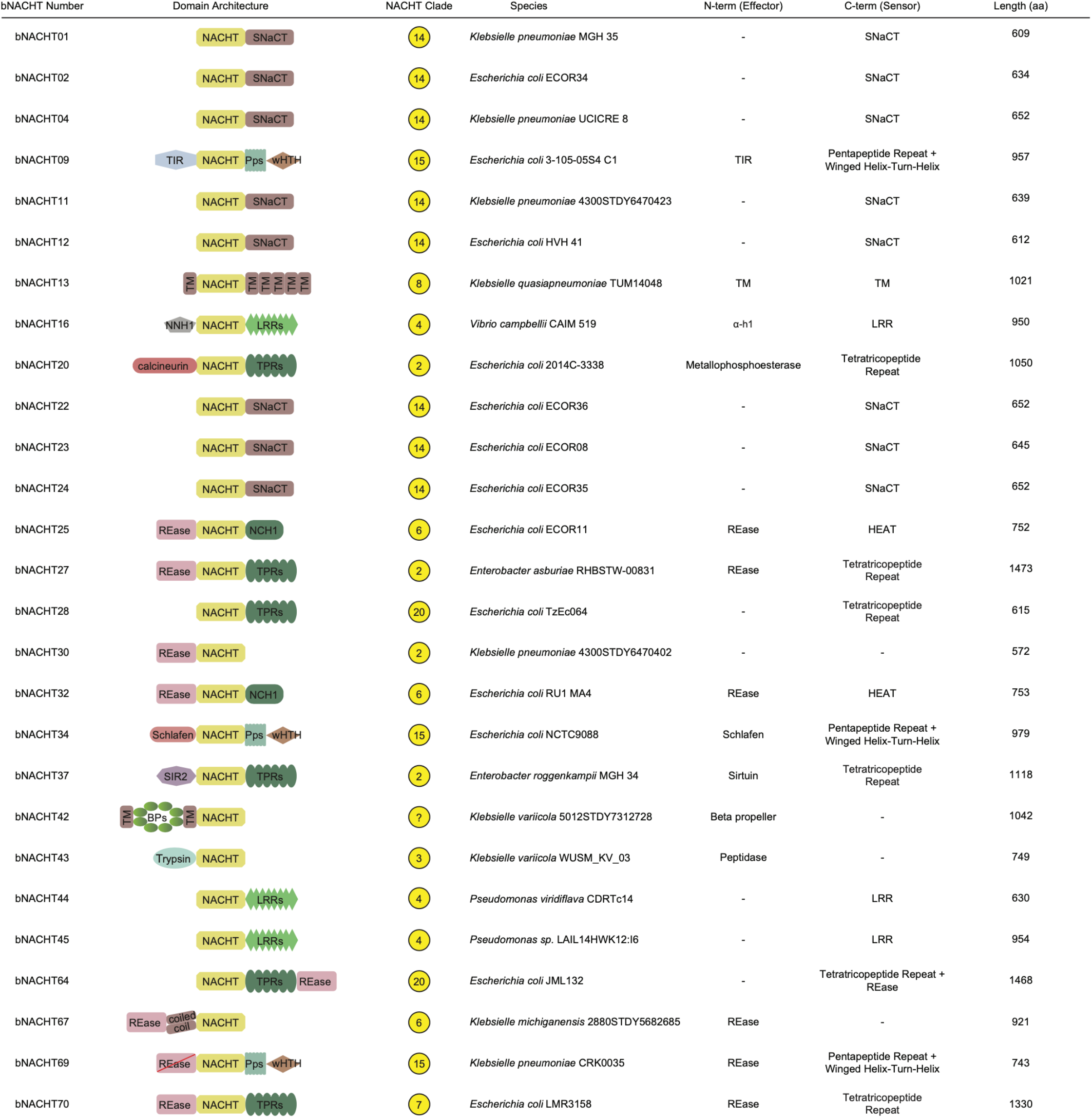
Domain architectures and details of experimentally tested NACHT proteins. Diagram of the N-terminal domains, central NACHT modules, and C-terminal domains of each experimentally tested bNACHT protein. The clade, species of origin, and length of each protein in amino acids (aa) are also indicated. Leucine rich repeat (LRR), Huntington, Elongation Factor 3, PR65/A, TOR (HEAT), Restriction endonuclease (REase), Toll/interleukin receptor (TIR), Transmembrane (TM), NACHT N-terminal helical domain 1 (NNH1), NACHT C-terminal helical domain 1 (NCH1).

**Figure S4.**
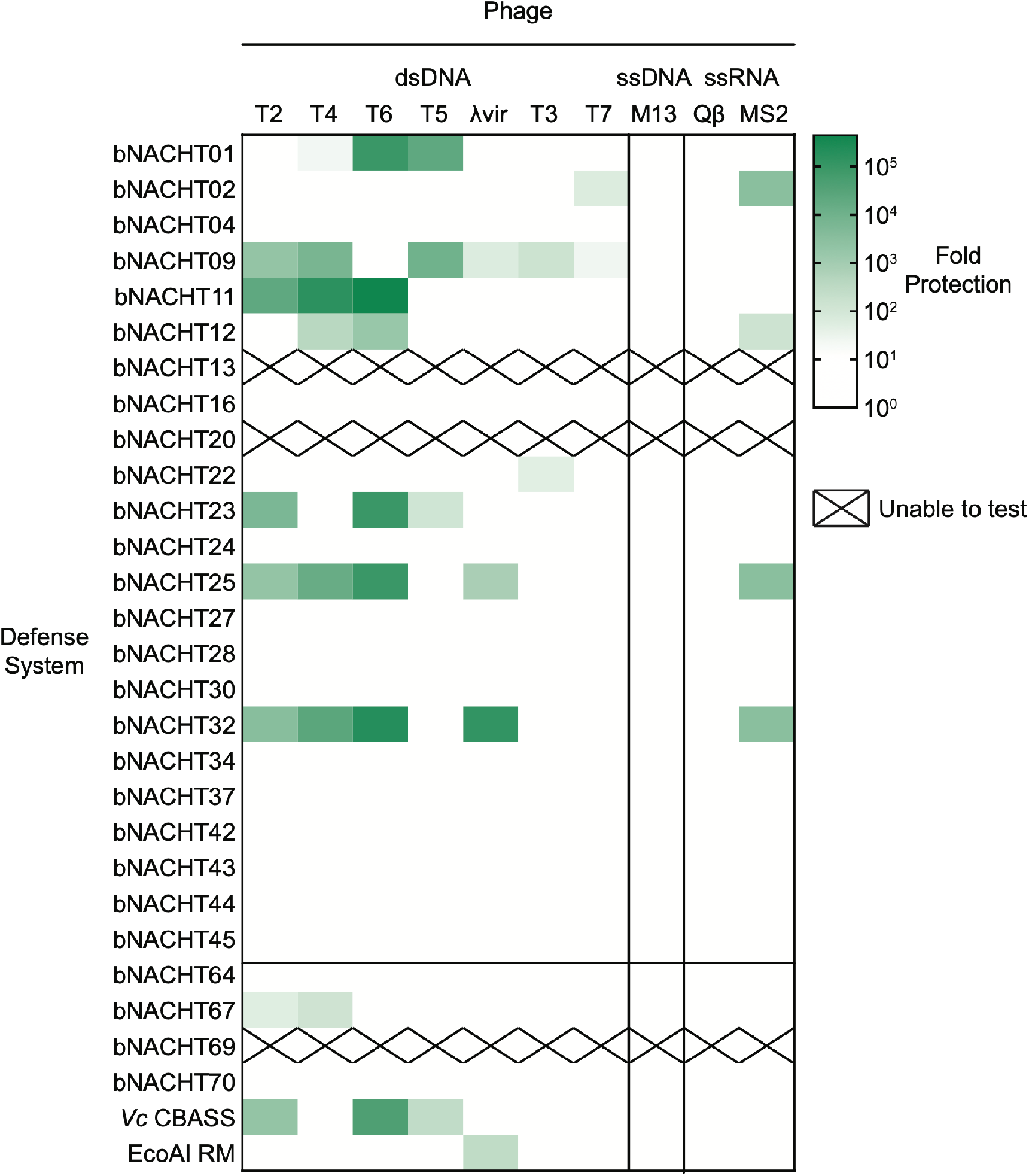
A screen for bacterial NACHT protein phage defense. Heat map of fold defense provided by the indicated bNACHT gene for a panel of diverse phages. Data are as described in **Figure 3** and show all experimentally interrogated bNACHT genes. Data were not collected for bacterial NACHT proteins that exhibited insufficient growth under these experimental conditions, indicated as “X”. Data represent the mean of *n* = *3* biological replicates. See **Table S2** and **Figure S3** for details on all 27 bNACHT genes analyzed. See **Figure S5** for raw efficiency of plating data.

**Figure S5.**
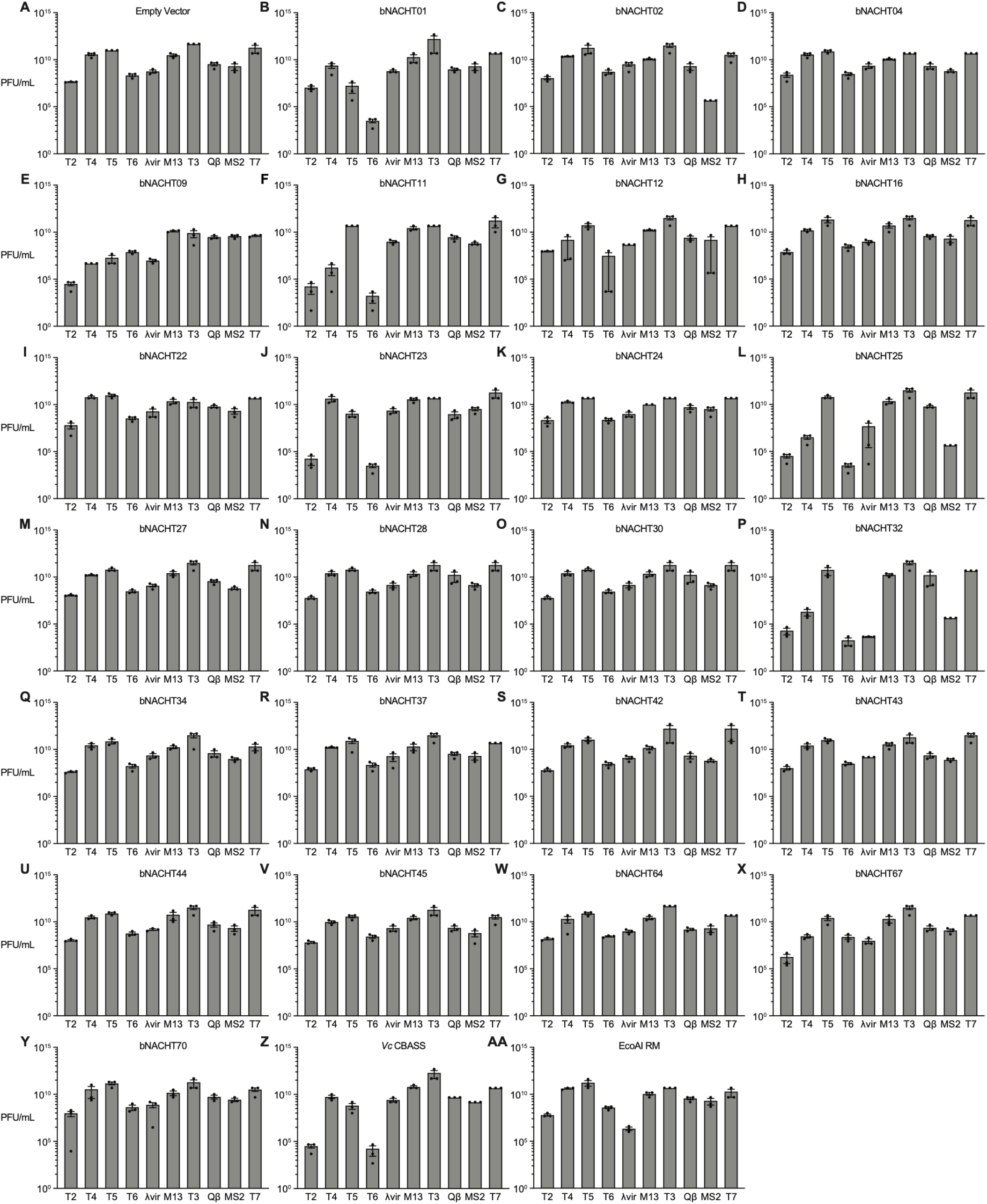
Efficiency of plating of phages infecting bacteria expressing NACHT proteins. Efficiency of plating of a phage panel infecting *E. coli* expressing the indicated bNACHT gene. The negative control is empty vector, which expresses an inactive *gfp* gene. Positive controls are *V. cholera* CBASS and *E. coli* UPEC-36 restriction modification system. Data represent the mean ± s.e.m. of *n = 3* biological replicates, indicated as individual points.

**Figure S6.**
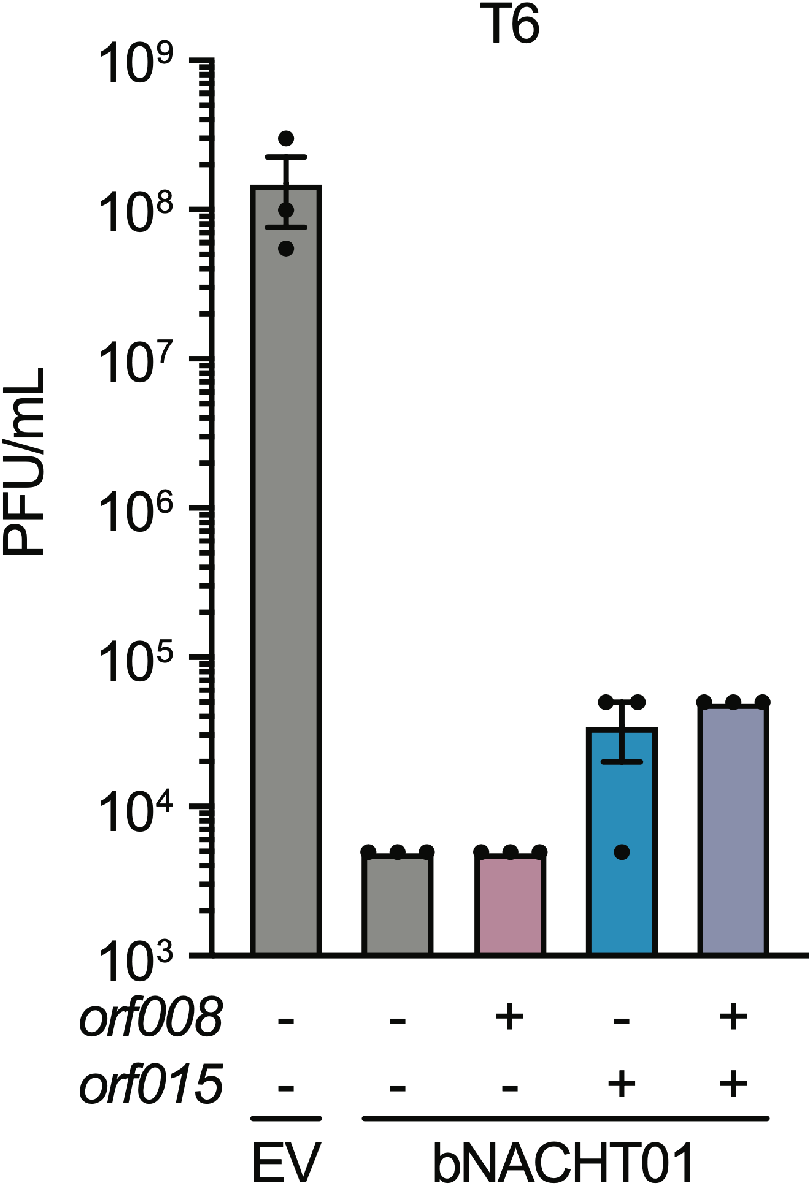
Effect of *orf008* and *orf015* expression on bNACHT01-mediated protection against phage T6. Efficiency of plating of phage T6 when infecting *E. coli* expressing bNACHT01 or empty vector (EV) on one plasmid and the indicated phage T5 gene(s) on a second plasmid. Expression of *orf008, orf015*, and *mCherry* is IPTG-inducible. (-) symbols denote induction of an *mCherry* negative control. (+) symbols denote induction of the indicated phage gene. Data represent the mean ± s.e.m. of *n* = *3* biological replicates, shown as individual points.

**Figure S7.**
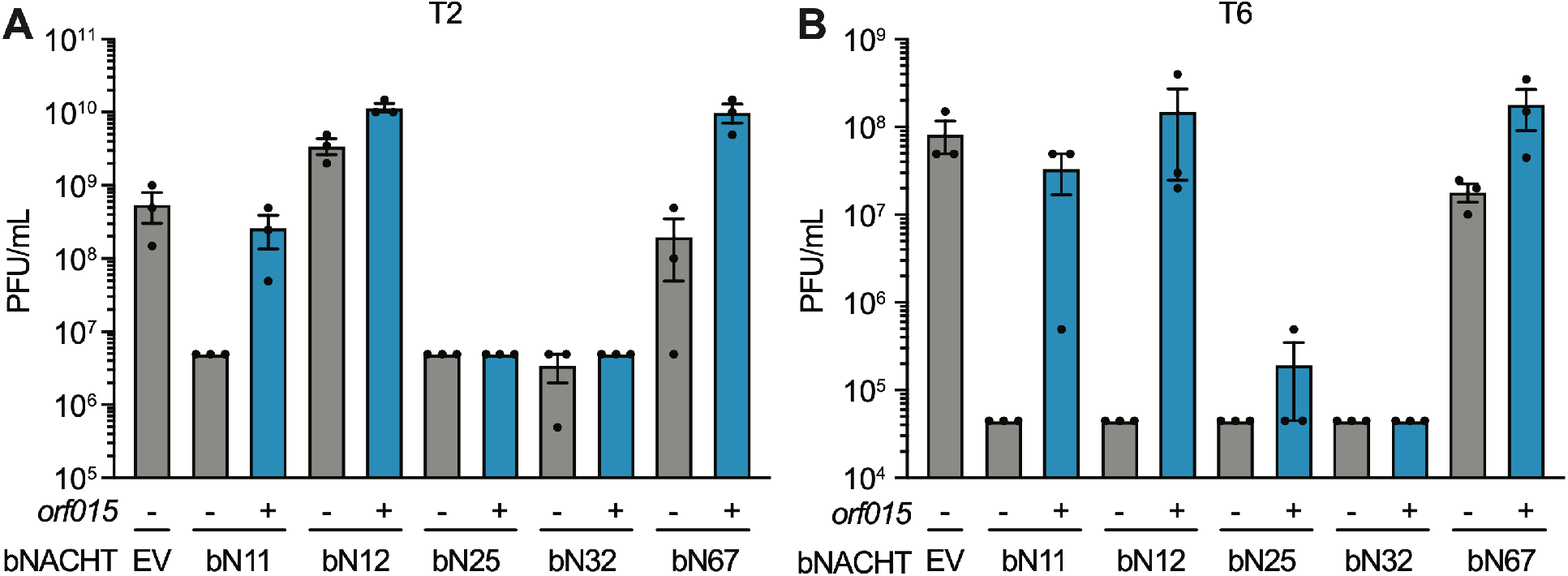
Effect of *orf015* expression on bacterial NACHT protein-mediated protection against phages T2 and T6. **(A)** Efficiency of plating of phage T2 when infecting *E. coli* expressing the indicated bNACHT gene on one plasmid and phage T5 *orf015* on a second plasmid. **(B)** Efficiency of plating of phage T6 when infecting *E. coli* expressing the indicated bNACHT gene on one plasmid and phage T5 *orf015* on a second plasmid. For A–B, expression of *orf015* or *mCherry* is IPTG-inducible. (-) symbols denote induction of an *mCherry* negative control. (+) symbols denote induction of *orf015*. Data represent the mean ± s.e.m. of *n* = *3* biological replicates, shown as individual points.

**Figure S8.**
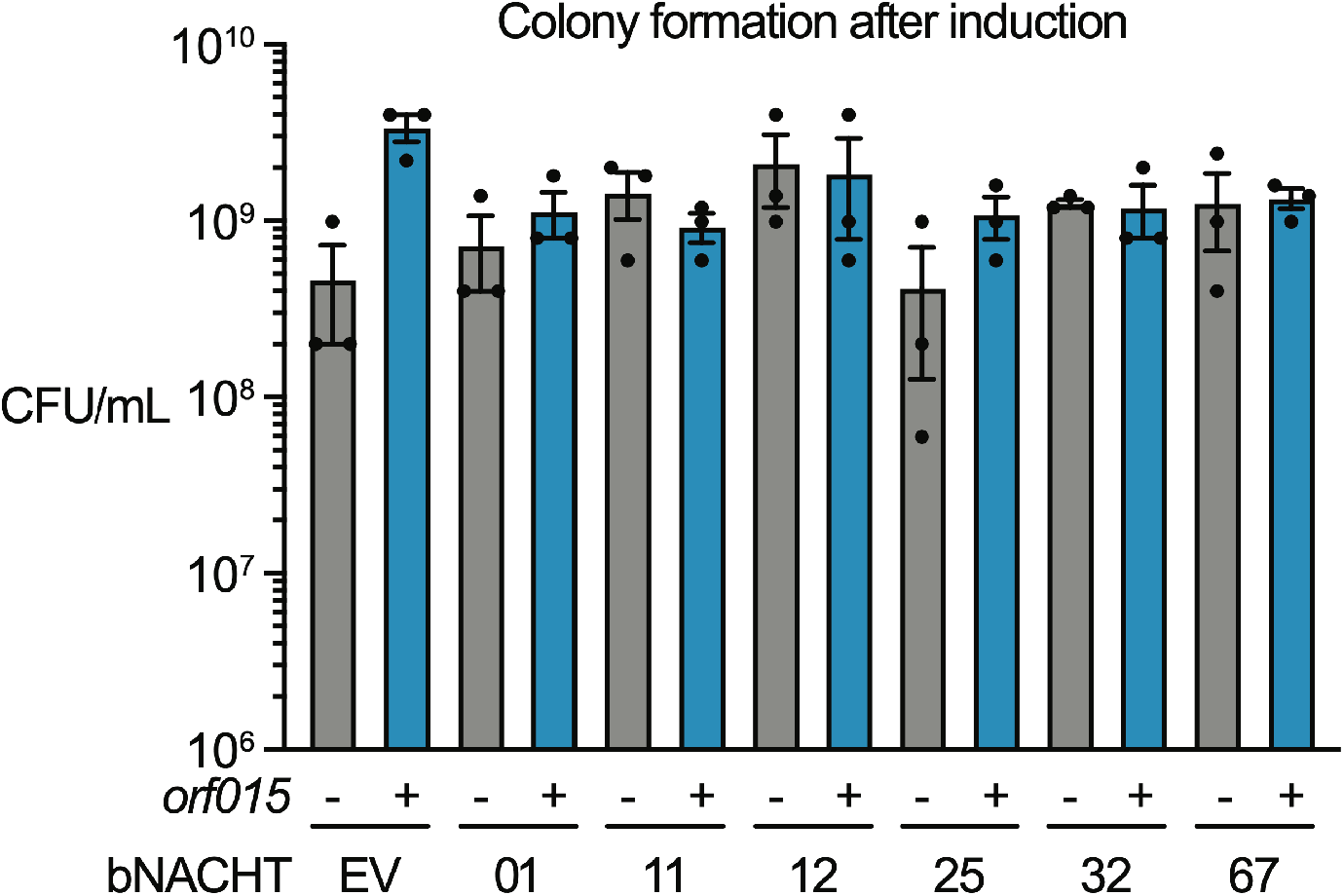
Effect of *orf015* expression on colony formation by bacteria expressing NACHT proteins. Quantification of colony formation of *E. coli* expressing the indicated bNACHT system on one plasmid and phage T5 *orf015* on a second plasmid. Expression of *orf015* or *mCherry* is IPTG-inducible. (-) symbols denote induction of an *mCherry* negative control. (+) symbols denote induction of *orf015*. Data represent the mean ± s.e.m. of *n* = *3* biological replicates, shown as individual points.

**Figure S9.**
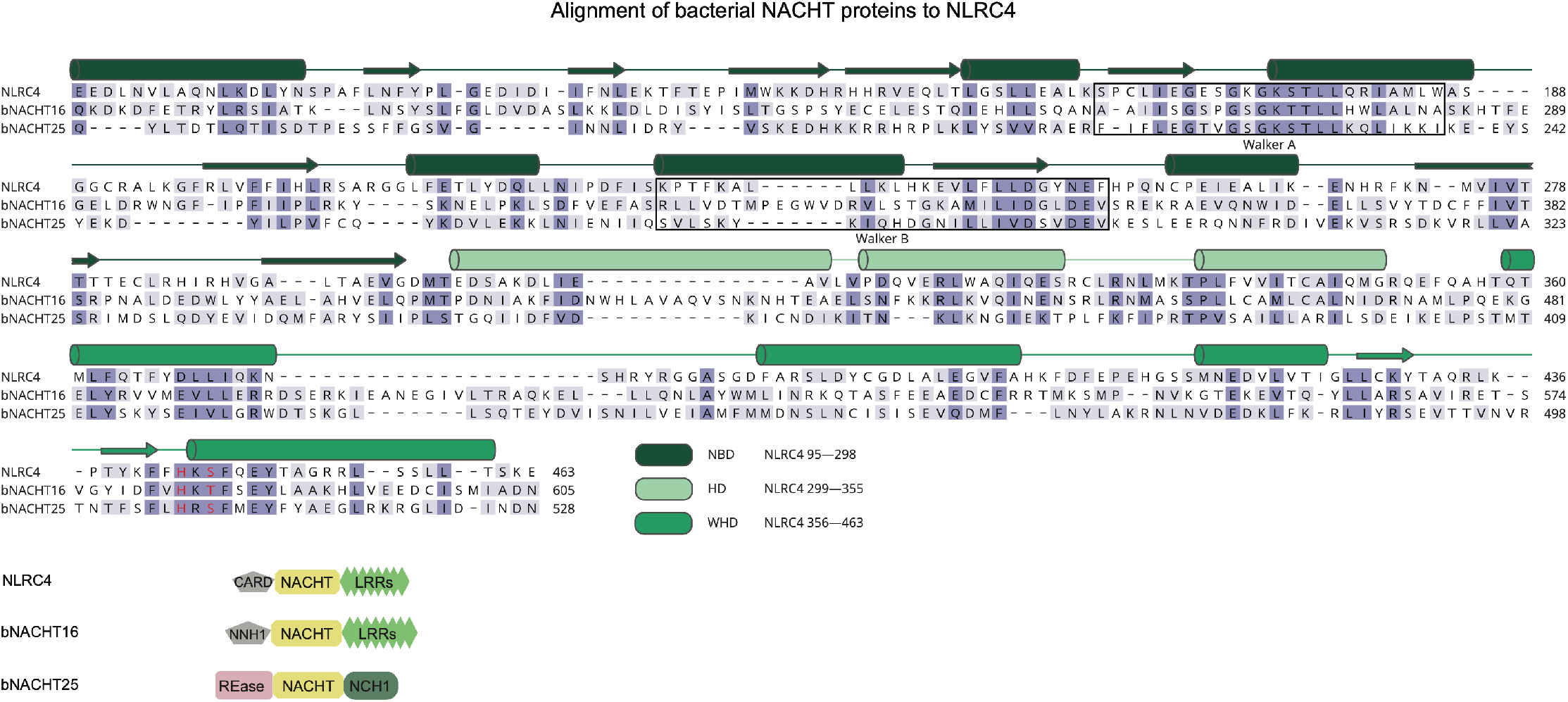
Protein alignment of NLRC4, bNACHT25, and bNACHT16. Protein alignment of the NACHT modules of NLRC4 (*Mus musculus;* NP_001028539), bNACHT16 (*Vibrio campbellii* CAIM 519; WP_005534681.1), and bNACHT25 (*E. coli* ECOR11; WP_001702659.1). The secondary structure of NLRC4 as determined by structure 4KXF in the PDB (Hu et al., 2013), is indicated above with alpha helices depicted as cylinders and beta sheets depicted as arrows. Secondary structure elements are color coded and labeled as in **Figure S1**. Amino acid residues are color-coded based on conservation in the sequence alignment. Black boxes indicate Walker A and B motifs. The conserved histidine and serine/threonine in the WHD is highlighted in red.

