## Supplementary material for "Bacterial NLR-related proteins protect against phage": Kibby et al. Table S3

Table S3. Rate of suppressor isolation for T5 infecting bNACHT01

| **Parent** | **Total PFU plated** | **Number of suppressor PFU** | **Rate of suppressor isolation (suppressor PFU per plated PFU)** |
| --- | --- | --- | --- |
| T5.1 | 8.1 × 10^7^ | 38 | 4.7 × 10^-7^ |
| T5.2 | 1.8 × 10^7^ | 15 | 8.3 × 10^-7^ |
| T5.3 | 1.7 × 10^7^ | 11 | 6.4 × 10^-7^ |
| T5.4 | 7.5 × 10^7^ | 7 | 9.3 × 10^-8^ |
