## Supplementary material for "Bacterial NLR-related proteins protect against phage": Kibby et al. Table S4

Table S4. T5 suppressor mutations in *orf008* and *orf015*

| **Phage parent** | **Suppressor phage** | ***orf008***^[1]^ **affecting mutation**^[2]^ | ***orf015***^[3]^ **affecting mutation** |
| --- | --- | --- | --- |
| T5.1 | T5.1.1 | Q73* | N4S |
| T5.1 | T5.1.2 | I16T | A64V |
| T5.2 | T5.2.1 | P17S | V69G |
| T5.2 | T5.2.2 | Promoter mutation (t4513c) | A64V |
| T5.3 | T5.3.1 | V64A | E22G |
| T5.3 | T5.3.2 | I16T | M1R |
| T5.3 | T5.3.3 | P17S | A64V |
| T5.4 | T5.4.1 | G56fs | F36fs |
| T5.4 | T5.4.2 | P17S | R62fs |
| T5.4 | T5.4.3 | I16T | ∆*orf009-012*, predicted promoter disrupting mutation |

1. The complete genome of bacteriophage T5 (ATCC11303-B) was used as a reference (NCBI genome accession AY587007.1). The NCBI protein accession number for the 83 amino acid protein Orf008 (hypothetical 9.2 kDa protein CDS) is AAX11945.1.
2. * indicates a premature stop codon and fs indicate a frame shift mutation.
3. The NCBI protein accession number for the 70 amino acid protein Orf015 is AAX11952.1.
