## Supplementary material for "Bacterial NLR-related proteins protect against phage": Kibby et al. Table S5

Table S5. All T5 suppressor mutations

| **Phage parent** | **Suppressor phage** | **Polymorphisms compared to parent**^[1]^ | **Polymorphism CDS**^[2]^ | **Predicted impact on CDS**^[3]^ |
| --- | --- | --- | --- | --- |
| T5.1 | T5.1.1 | g4261a | *orf008* | Q73* |
|  |  | a8017g | *orf015* | N4S |
| T5.1 | T5.1.2 | a4434g | *orf008* | I16T |
|  |  | c8197t | *orf015* | A64V |
| T5.2 | T5.2.1 | g4432a | *orf008* | P17S |
|  |  | t8212g | *orf015* | V69G |
| T5.2 | T5.2.2 | t4513t | Intergenic | *orf008* promoter mutation |
|  |  | c8197t | *orf015* | A64V |
| T5.3 | T5.3.1 | a4290g | *orf008* | V64A |
|  |  | a8071g | *orf015* | E22G |
| T5.3 | T5.3.2 | a4434g | *orf008* | I16T |
|  |  | t8008g | *orf015* | M1R |
| T5.3 | T5.3.3 | g4432a | *orf008* | P17S |
|  |  | c8197t | *orf015* | A64V |
| T5.4 | T5.4.1 | ∆t4313 | *orf008* | G56fs |
|  |  | t8112tt | *orf015* | F36fs |
| T5.4 | T5.4.2 | g4432a | *orf008* | P17S |
|  |  | ∆t8190 | *orf015* | R62fs |
| T5.4 | T5.4.3 | a4434g | *orf008* | I16T |
|  |  | ∆4794–7468 | *orf009–orf012* | ∆*orf009–012,* alters *orf015* promoter |

1. Phage T5 was obtained from the CGSC (CGSC 12144) and sequenced genomes were compared to NCBI genome accession AY587007.1. Polymorphisms found in the parent genomes were disregarded.
2. Orf008 is an 83 amino acid protein whose NCBI protein accession is AAX11945.1. Orf015 is a 70 amino acid protein whose NCBI protein accession is AAX11952.1.
3. * indicates a premature stop codon and fs indicate a frame shift mutation.
